# Caregiver-infant interactions selectively shape emerging functional connectivity in the neonatal brain

**DOI:** 10.64898/2026.05.13.724776

**Authors:** Laura Carnevali, Borja Blanco, Maria Rozhko, Matthew Weatherhead, Mark H. Johnson, Sarah Lloyd-Fox, The PIPKIN Study Team

## Abstract

From birth, human infants engage in multi-modal social exchanges with caregivers that involve the coordination of gaze and touch to guide attention and support neurodevelopment. However, little is known about the association between these first interactive experiences and the functional organization of the developing brain during the first postnatal month, a window of remarkable brain growth in humans. We address this gap by combining microanalytic coding of caregiver– infant interactions with task-free functional connectivity (FC), measured using high-density diffuse optical tomography (HD-DOT) in infants’ homes during the first postnatal month. Task-free FC measures the intrinsic functional organization of the developing brain, shedding light on the early development of neural systems supporting perception, regulation, and social interactions. Infants were assessed up to three times (1 week, 2 weeks, 1 month), enabling characterization of both early FC and its rapid developmental change. Caregiver-infant interactions were associated with both concurrent organization and rapid longitudinal change in FC. Dyadic engagement in the context of face-to-face interaction was associated with the refinement of short-range connectivity and the integration of long-range connectivity particularly between social brain regions, while affectionate touch was associated with general increases in long-distance connectivity. These results demonstrate that caregiving experiences influence the development of the brain’s functional architecture in the first postnatal month, highlighting a critical window for shaping infant brain function.

**Significance Statement:** The first postnatal weeks are a period of rapid brain development, when the brain may be especially sensitive to experience. A key early experience is social interaction between infants and their caregivers, yet it remains unknown if these interactions influence neonate brain function. By combining observations of caregiver–infant interactions at one month with repeated home-based brain imaging, we show that variations in dyadic engagement and affectionate touch influence how infant brain functional connectivity is organized and develops. These findings highlight the importance of supporting perinatal life and early interactions for infant brain development.

## Introduction

Infancy represents a period of very rapid brain development, during which cortical circuits rapidly specialize, potentially through experience-dependent processes (1–5). From birth, the human brain is organized for efficient communication within local functional networks and for global integration via selective long-range connections (6). Functional connectivity (FC) measures the temporal coordination of activity between brain regions, providing a window into how early developing local networks and high-order long-range networks communicate to support perception, cognition, and social behaviour (7). Primary sensorimotor networks, including visual and auditory systems, consolidate early on in development to support local perception and action (5, 8–12), while association, high-order, long-range networks including default mode, frontoparietal, and attention networks display rapid reorganization and high individual variability within the first postnatal months (5, 13, 14). Understanding this individual variability is critical as these networks underlie processes that are essential for social cognition; including face processing, joint attention, and social learning (14–19).

Over the neonatal period, one of the main experiences that forms the basis of infant learning are social interactions. From birth, infants continuously engage in multisensory caregiver-infant exchanges, in which gaze, touch, and vocalizations direct attention and guide the refinement of cortical networks (20–23). These ostensive cues signal communicative intent, orient the infant to relevant environmental stimuli, and provide a foundation for early social learning (24, 25). Eye contact is particularly salient, amplifying attention to faces, joint attention, and social engagement (26–29). Touch is the first sensory modality to develop in utero and affectionate touch, in particular, is associated with arousal regulation, stress reduction and attention toward socially relevant cues, while also promoting affiliative bonds and social learning (30–33). Caregivers use touch and gaze contingently in naturalistic day-to-day interactions (34, 35). Yet, to date, no study has examined how these early interactive experiences relate to functional brain organization in the first postnatal months (36).

FC is sensitive to both maturational processes and early experiences (6, 37–39). Variability in FC patterns has been linked to social attention, behavioural regulation, and risk for later social difficulties (14, 16). These associations are particularly pronounced within the ‘social brain’ a set of regions including the superior temporal sulcus (STS), temporoparietal junction (TPJ), and parts of the prefrontal cortex (PFC), which detect and interpret social signals (40). Despite evidence that regions within these networks show increasingly specialized responses to social stimuli in the first year of life (41–43), very little is known about how real-life, moment-to-moment caregiver-infant interactions shape functional brain organization in the earliest postnatal weeks (36). Such interactions provide a rich, multisensory environment that scaffolds social attention, regulation, and learning, and may contribute to functional network organization during a highly plastic developmental window. Understanding how these early social interactions influence network organization is therefore essential for identifying the mechanisms by which early social experience shapes functional brain development (5, 44).

Individual differences in FC have been linked to features of the caregiving environment, highlighting early influences on brain network organisation. Neonatal connectivity varies with caregiver factors such as maternal stress and mental health, particularly in frontoparietal and ventral attention networks (45, 46), and with infant characteristics such as behavioural temperament at one month, which predicts FC in frontoparietal, and default mode networks linked to regulation and orienting (47). Early experiences continue to shape FC across development: maternal sensitivity at five months positively associates with default mode network connectivity (48) and maternal touch frequency at five years predicts posterior STS–dorsal mPFC connectivity (49). Together, these findings suggest that a more enriched and responsive caregiving environment may support functional network development. At the same time, they indicate that early functional network specialisation is not determined by the infant or caregiver alone, but by their dynamic, reciprocal interactions, exchanges of attention, touch, and affect that together help shape the developing brain.

Most prior work relies on laboratory-based tasks, parental reports, or measurements taken months later, and very few studies combine functional connectivity measures with observations of naturalistic caregiver-infant interactions. As a result, the earliest interactive experiences between infants and caregivers remain unexplored in relation to functional network development. Here, we address this gap by combining microanalytic coding of naturalistic caregiver-infant interactions in the home with task-free FC data acquired via high-density diffuse optical tomography (HD-DOT) during the first month of life. As part of the PIPKIN study (50), infants were visited at home up to three times (approximately at one week, two weeks, and at 1 month), enabling us to capture both early FC organization and its short-term developmental dynamics. This dense longitudinal sampling targets the neonatal period, allowing us to examine how functional brain networks change over a window characterized by exceptionally rapid neural reorganization (6).

This study provides a unique opportunity to link naturalistic caregiver-infant interactions to both cross-sectional and longitudinal changes in functional networks during the first postnatal month. Prior work in older infants and children suggests that enriched caregiving is associated with stronger long-range connectivity, particularly within social brain regions. We therefore hypothesize that higher levels of dyadic engagement and more frequent affectionate touch will be associated with stronger functional connectivity, particularly within the social brain. By focusing on this early time window, our study sheds light into how interactive social experiences shape the developing functional architecture of the human brain, providing insights with scientific and societal relevance, including implications for supporting optimal parent-infant interactions.

## Results

To examine how early caregiver–infant interaction behaviours relate to functional connectivity (FC) during the first postnatal month, we first assessed whether dyadic engagement and affectionate touch during naturalistic interactions at one month of age were associated with the strength (Fisher-z transformed correlations) and extent (number of connections) of HbO FC (Figure 1) across brain regions and distances (HbR results are included in supplementary materials). Dyadic engagement measures the interaction between parent and child, rather than capturing the behaviour of either participant in isolation. Here, it was measured using a combined index of caregiver-to-infant gaze, infant-to-caregiver gaze, and touch occurring while the infant was looking at the parent. We also examined affectionate touch, defined as gentle, positive tactile behaviours (e.g., caressing, handholding, kissing, and playful touch), which represent a key component of early caregiver-infant communication and is thought to play an important role in regulating infant arousal and supporting socio-emotional development. We then examined longitudinal changes in FC across the first month of life, testing whether these changes differed by seed region and connection distance – short-(≤30 mm), medium-(30–60 mm), or long-range (>60 mm), following Wang et al. (2021) (51) – and whether individual differences in these changes were associated with caregiver–infant interaction measures.

**Figure 1.**
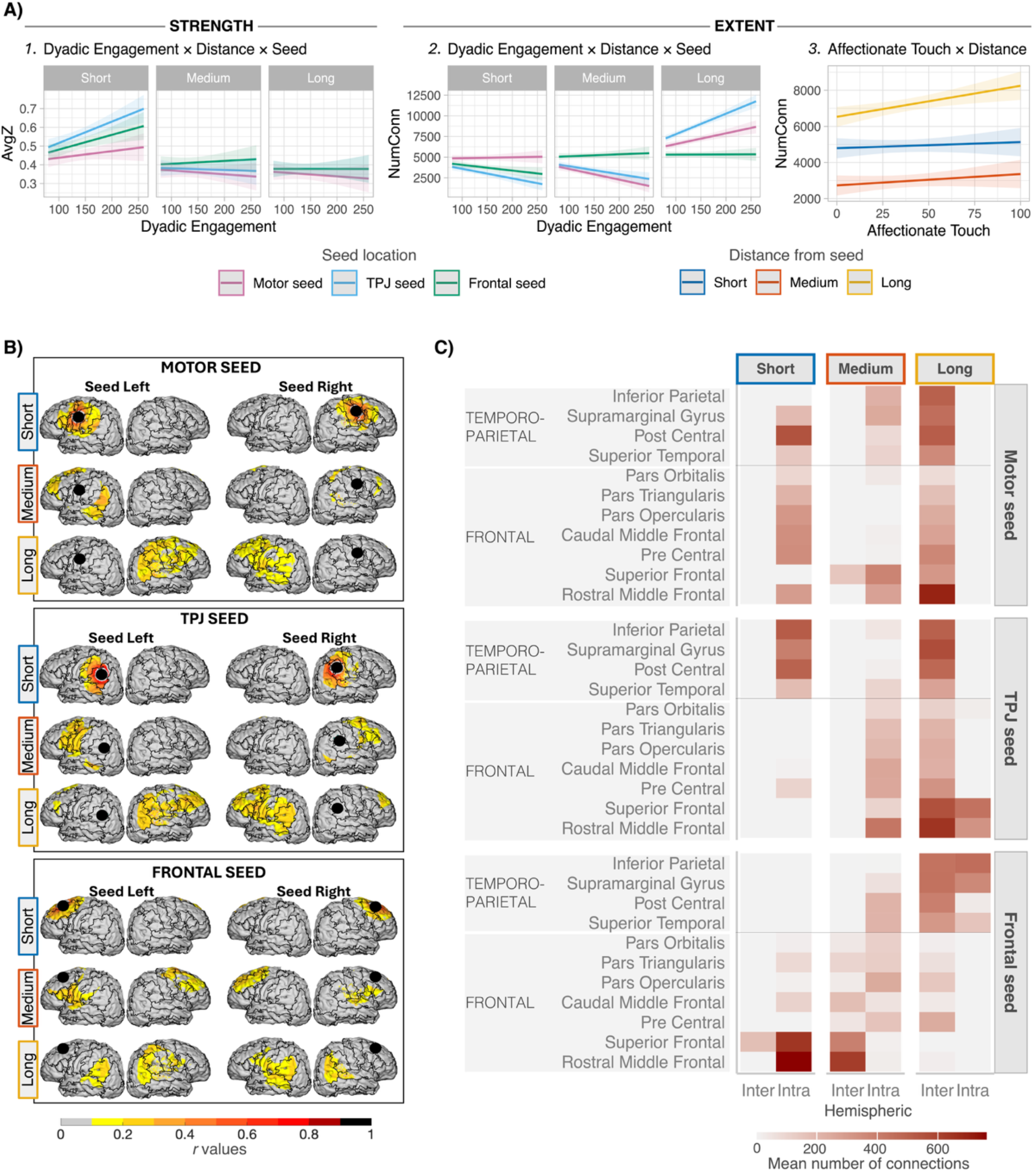
1-month HbO functional connectivity and caregiver-infant interaction associations. Functional connectivity was quantified in terms of strength (average Fisher-z transformed correlations, labelled “AvgZ”) and extent (number of connections, labelled “NumConn”). Parent-infant interaction measures included Dyadic Engagement, defined as a composite proportion index computed as the sum of the percentages of infant-to-parent gaze, parent-to-infant gaze, and affectionate touch occurring during periods of infant social attention (i.e., when the infant was looking at the parent), with each component expressed relative to the total interaction time. This approach preserves variability across modalities (gaze and touch) rather than averaging them. Affectionate Touch was defined as the percentage of total interaction time during which caress-like, playful, kissing, and hand-holding behaviours were observed. In both cases, higher values indicate greater relative behavioural occurrence. **(A)** Effects visualisations for the interaction of Dyadic Engagement, Distance and Seed on HbO Strength (A1, average Fisher-z transformed correlations) and Extent (A2, number of connections), as well as for Affectionate Touch and Distance on Extent (A3). The same effects emerged as significant for HbR, as reported in the Supplementary materials. **(B)** Group-level maps of functional connectivity across seeds and hemispheres, at different distances. Connectivity values (*r* values) represent Pearson correlation coefficients between haemodynamic signals, with higher values indicating stronger functional connectivity. Seeds are indicated by black dots. **(C)** Group-level heatmaps showing the average number of connections from each seed to each of the anatomical parcels within and across hemispheres.

### 1.1 Caregiver–infant interactions shape functional connectivity at one month

Dyadic engagement was related to stronger (higher Fisher-z transformed correlations) but fewer short-range connections for TPJ and frontal seeds, alongside reduced medium-range and increased long-range connectivity extent (number of connections) for TPJ and motor seeds (Figure 1A). For short-range connections, dyadic engagement was associated with increased FC strength, with significant positive effects in the TPJ (β = 1.14 × 10^−3^, SE = 2.87 × 10^−4^, t(47.66) = 3.97, p < .001) and frontal seeds (β = 7.87 × 10^−4^, SE = 2.87 × 10^−4^, t(47.66) = 2.74, p = 0.009). For short-range connectivity extent, dyadic engagement showed a negative association for TPJ (β = −11.6, SE = 2.76, t(65.2) = −4.21, p < .001) and frontal seeds (β = −6.9, SE = 2.76, t(65.2) = −2.5, p = 0.015), indicating fewer short-range connections at higher dyadic engagement scores. Short-range connections from the TPJ seed were mostly targeting intra-hemispheric temporoparietal and precentral regions (Figure 1B, C). Those from the frontal seed were primarily intra-hemispheric within the frontal lobe, with additional inter-hemispheric links to superior frontal regions (Figure 1B, C). For medium-range connections, dyadic engagement was associated with reduced connectivity extent for the motor (β = −12.9, SE = 2.76, t(65.2) = −4.67, p < .001) and TPJ (β = −9.65, SE = 2.76, t(65.2) = −3.49, p < .001) seeds. These connections were mostly intra-hemispheric, extending to temporal and posterior frontal regions from TPJ, as well as posterior-frontal and temporoparietal from the motor seed (Figure 1B, C). In contrast, for long-range connections, dyadic engagement was associated with increased connectivity extent, particularly for TPJ (β = 24.9, SE = 2.76, t(65.2) = 9.01, p < .001) and motor seeds (β = 12.99, SE = 2.76, t(65.2) = 4.7, p < .001). For TPJ, increase in long-range connections included intra-hemispheric connections to anterior frontal regions and inter-hemispheric links to frontal lobe more broadly (Figure 1B, C). No significant effects emerged in connectivity strength for medium- and long-range connections.

We next examined the effect of affectionate touch on FC and found that it was positively associated with the number of long-range connections across seeds (β = 17.2, SE = 5.81, t(95.32) = 2.96, p = 0.004), with no significant effects on short- or medium-range connectivity or connectivity strength (Figure 1A). The same effects emerged as significant for HbR, as reported in the Supplementary materials.

Notably, while previous works have predominantly examined associations between early behaviours and connectivity strength, our model fit was higher when predicting extent (conditional *R*^2^ = 79%, marginal *R*^2^ = 74%) than strength (conditional *R*^2^ = 66%, marginal *R*^2^ = 39%), suggesting that extent was a more sensitive measure in this sample. Model comparisons for strength and extent and fit indices are reported in Supplementary Table 4.

### 1.2 Longitudinal changes in functional connectivity across the first month

To examine the rapid postnatal reorganization of functional connectivity, we assessed longitudinal change across the first postnatal month by modelling within-subject variation in connectivity extent. Developmental change was quantified as the individual-level slope of the number of suprathreshold Fisher-z-transformed connections as a function of gestational age (weeks post-conception), allowing participant-specific trajectories to be estimated across distances and hemispheres.

We found that longitudinal change in connectivity extent was significantly predicted by the Distance × Seed interaction (F(4, 288) = 13.25, p < .001) (Figure 2 A). Over the first month, long-distance connections increased overall across all seeds (Motor: β = 532, 95% CI [299, 765], p < .001; TPJ: β = 542, 95% CI [309, 775], p < .001; Frontal: β = 543, 95% CI [310, 776], p < .001). Short- and medium-distance connectivity showed seed-specific developmental patterns for the motor and TPJ seeds. The motor seed exhibited increased short-range and decreased medium-range connectivity (short: β = 183, 95% CI [−50, 416], p = .032; medium: β = −224, 95% CI [−457, 9], p = .012); whereas the TPJ seed showed decreased short-range and increased medium-range connectivity (short: β = −167, 95% CI [−400, 66], p = .045; medium: β = 240, 95% CI [7, 473], p = .008). Developmental trajectories of connectivity extent showed considerable inter-individual variability (Figure 2B).

**Figure 2.**
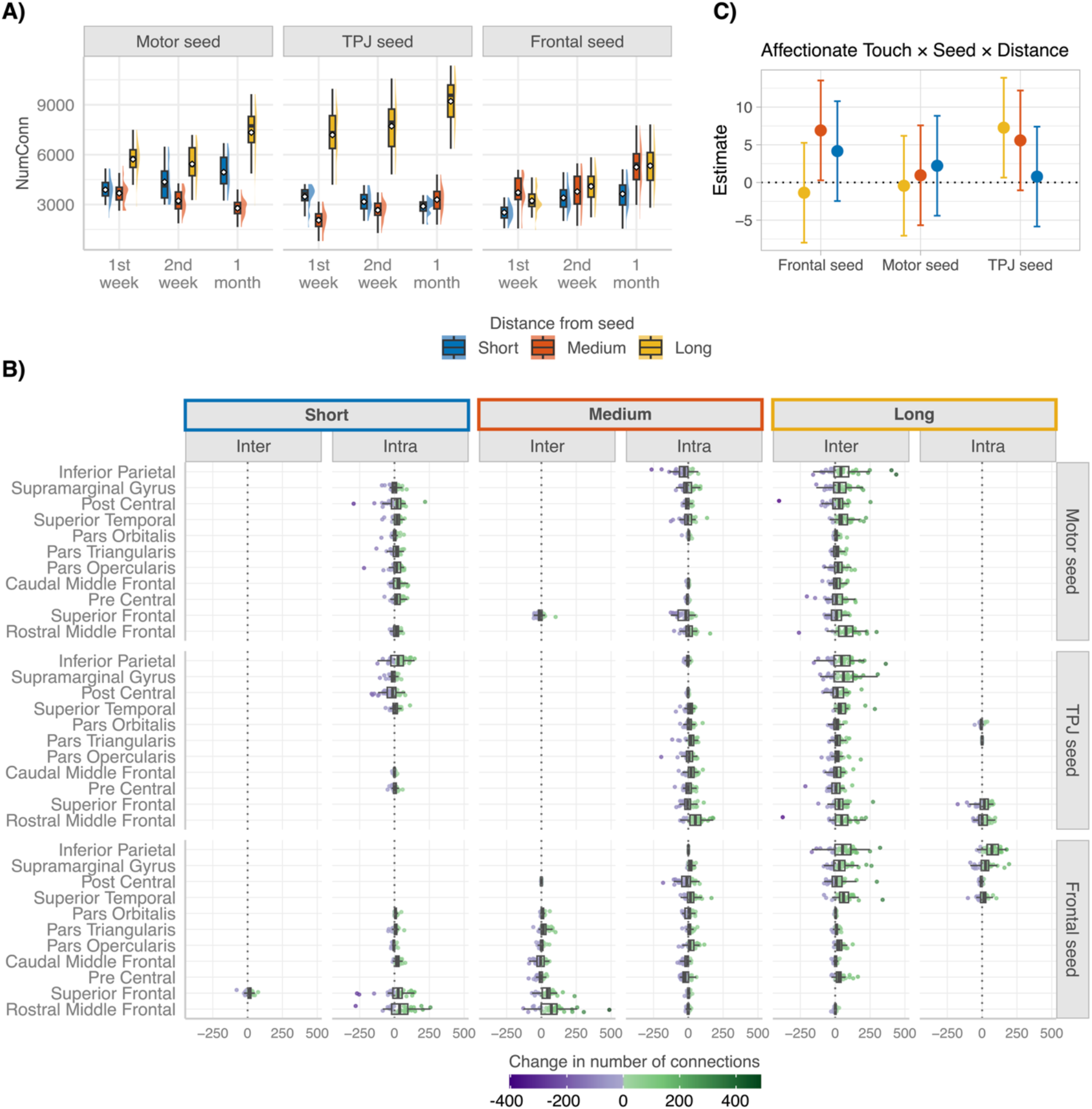
Longitudinal change in HbO connectivity extent. **(A)** Distributions of number of connections for each distance within specific seed placements, across timepoints (first week, second week and one month postnatal). **(B)** Individual slopes (number of connections ~ age in weeks post-conception) quantifying developmental change in the number of connections between each seed and each of the parcels, across distances and hemispheres. Individual points represent participants. **(C)** β estimates (± 95% CI) from the Affectionate Touch × Distance × Seed interaction term, predicting longitudinal change (individual slopes) of FC extent over time across combinations of Seed and Distance. Positive estimates indicate that higher affectionate touch was associated with steeper longitudinal increases in FC extent.

Individual differences in longitudinal change of FC extent were associated with parent tactile behaviour during early interactions. At longer distances, higher levels of affectionate touch were associated with increased longitudinal change in FC, with seed-specific effects emerging in the interaction model (Figure 2C). Specifically, the Affectionate Touch × Distance × Seed interaction significantly predicted longitudinal change (slopes) in connectivity extent (Frontal: medium-range β = 6.90, SE = 3.38, t(143.85) = 2.04, p = .043; TPJ: long-range β = 7.27, SE = 3.38, t(143.85) = 2.15, p = .033), whereas no significant effects were observed for short-range connectivity across seeds. These effects indicate that greater affectionate touch was specifically associated with steeper increases in long-range connectivity for TPJ, and in medium-range connectivity for frontal systems (Figure 2C). For dyadic engagement, no significant associations were observed with longitudinal changes in connectivity extent across any distance or seed region. Model comparisons and fit indices are reported in Supplementary Table 4.

## Discussion

Our findings provide evidence that early caregiver-infant interactions are associated with both the organization and rapid developmental change of brain functional connectivity (FC) during the first postnatal month. We linked naturalistic caregiver-infant behaviours within the first month postnatal to both cross-sectional and longitudinal changes in functional brain networks. Notably, we show that distinct dimensions of early caregiving differentially influence the developing neonatal brain: dyadic engagement promotes local refinement of emerging circuits and their long-range integration, particularly in regions supporting social and affective processing, whereas affectionate touch associated with long-range network integration at a broader scale. These complementary influences may reflect distinct underlying mechanisms. Dyadic engagement is inherently reciprocal and may reinforce specific networks through mutual attunement and temporally contingent social exchange, selectively modulating circuits that are still emerging. In contrast, affectionate touch is largely caregiver-driven and present from birth or earlier, providing a stable source of multisensory and affective input that may broadly facilitate large-scale network integration. This interpretation aligns with prior work linking tactile and affective caregiving to neural systems involved in social processing, including increased activity and connectivity in mentalizing-related regions (49) and enhanced caregiver-infant synchrony (52, 53). Within this framework, we first consider each caregiving dimension and then examine how these effects relate to network-specific developmental trajectories.

Dyadic engagement during face-to-face interactions was associated with stronger but fewer short-range connections, alongside increased long-range connectivity. Notably, effects on connectivity strength were restricted to short distances and observed in TPJ and frontal networks, but not in the motor network. Given that short-range connectivity dominates neonatal brain organization (54, 55) and that sensorimotor networks develop earlier and more rapidly than higher-order networks (5, 56, 57), these findings suggest that dyadic engagement selectively modulates the strength of locally organized connections in networks that are still emerging. At the same time, the reduction in connectivity extent at short ranges may be consistent with a process of selective stabilisation, whereby some connections are strengthened while others are weakened or pruned. Developmental changes in FC at longer distances were primarily reflected in connection extent rather than strength, suggesting that the neonatal brain initially prioritizes establishing new links, while selective strengthening may emerge later. This is broadly consistent with models in which early development is characterised by exuberant formation of connections, followed by activity-dependent pruning and increasing network selectivity (58). However, we did not observe a reduction in FC for long distances, possibly because of the early time window of our study, but we would expect such effects at later developmental timepoints. These region- and distance-specific effects provide evidence that early social experience influences network-specific trajectories to shape emerging functional organization. Importantly, supplementary analyses confirmed that longitudinal changes in FC did not explain a substantial amount of variance in the dyadic engagement composite score, indicating that early social interactions shape functional connectivity, and not the opposite (Supplementary Table 7).

Affectionate touch showed a different profile. At one month, higher levels of caregiver-provided touch were associated with a greater number of long-distance connections across seed regions, without evidence of selective effects across networks. Longitudinally, affectionate touch associated with changes in connectivity extent, particularly within temporoparietal and frontal regions. These regions are central to emerging social and affective processing (42, 59, 60), suggesting that caregiver-provided touch may facilitate rapid network reconfiguration during a critical developmental window by supporting the formation of distributed functional connections. While direct evidence linking touch to FC in neonates remains limited, converging work in older children supports the plausibility of these effects: in 5-years-olds, higher frequency of maternal touch predicts stronger resting-state functional connectivity between right pSTS/TPJ and a distributed social-brain network, including bilateral temporal and frontal cortices, insula, and dorsal mPFC (49). Skin-to-skin contact in preterm infants is similarly associated with accelerated hemispheric maturation and enhanced functional connectivity (61, 62). Mechanistically, affectionate touch engages somatosensory, temporal, and frontal circuits, modulates regulatory and autonomic systems, and promotes caregiver–infant neural synchrony, providing a plausible pathway through which repeated touch shapes long-range functional connectivity (35, 63).

The above is interpretable in the context of network-specific developmental trajectories in early life. Our findings suggest that early FC changes initially reflect the formation of new functional connections rather than the selective strengthening of already established links. TPJ connections appear to be in a phase of spatial expansion, increasing their extent as they integrate into distributed systems. Frontal networks show broad early recruitment, gradually establishing nascent connections that may later specialize and strengthen. Motor networks, already relatively developed, primarily undergo local modularization and show little sensitivity to variability related to early interaction patterns. This seed- and distance-specific pacing aligns with broader posterior-to-anterior and primary-to-higher-order sequences of early network development (6, 44, 54, 64), providing a framework for interpreting how early social experiences interact with intrinsic developmental trajectories to shape emerging functional brain organization.

Temporoparietal regions exhibited a distance- and time-dependent profile, which was susceptible to interactional influences. At one month, the TPJ showed local connectivity within temporoparietal areas alongside medium- and long-range connections to frontal regions, consistent with the early emergence of networks implicated in social processing (60). Longitudinally, TPJ connectivity shifted away from short-range connections toward increased medium- and long-range connectivity, reflecting progressive integration with distributed networks. This pattern aligns with previous work suggesting that neonatal FC is initially dominated by short-range connections, particularly between anatomically adjacent regions (65, 66), while long-range connectivity emerges more gradually, undergoing substantial reorganization even within the first postnatal months (5). This view is also supported by the developmental trajectories we observed in the frontal network. At one-month, the frontal seed exhibited predominantly local connectivity at short and medium distances, with long-range links mainly to temporoparietal regions. Longitudinally, frontal connectivity increased broadly across distances, albeit without the distance-dependent differentiation observed for TPJ and motor networks. This may reflect an initial, diffuse recruitment phase in which frontal networks establish functional connections across the brain rather than selectively tuning specific links, consistent with the protracted development of frontal systems reported in prior work (5). Nonetheless, correlations between short- and medium-range frontal connectivity and affectionate touch are consistent with the view of frontal network being particularly sensitive to environmental caregiving context (46).

Sensorimotor regions are known to exhibit robust early connectivity, with strong intra- and inter-hemispheric coupling, reflecting their foundational role in supporting early sensorimotor functions (67, 68). Consistent with this, motor regions showed longitudinal increases in short-range and decreases in medium-range connectivity, reflecting local specialization and modularization within the sensorimotor cortex (69). In parallel, long-range motor connectivity increased over time, largely driven by widespread interhemispheric links rather than being restricted to homologous motor regions. These patterns align with evidence that neonatal somatosensory cortex acts as an early hub, characterized by high centrality, dense local connectivity, and widespread links to distributed regions, supporting both somatotopic specialization and global integration across networks (58). Notably, these developmental changes in motor networks were largely unrelated to dyadic engagement or affectionate touch, suggesting limited influence of early interactions patterns on networks that are already relatively developed in early perinatal life, such as the sensorimotor network (6).

Taken together, these results support a model in which early social experience biases how some developing brain networks negotiate the balance between functional segregation and integration during a period of rapid change. Dyadic engagement contributes to local refinement within short-range networks, whereas affectionate touch may scaffold global expansion of long-distance connectivity as large-scale networks emerge. Within the first postnatal month, when long-range connectivity increases robustly, such experiential influences may be particularly consequential, subtly shaping functional organisation rather than simply adding or removing connections. Crucially, sensitive caregiving, environmental enrichment, and social support have been shown to buffer the effects of prenatal stress and socioeconomic disadvantage on neural development and later neurocognitive outcomes (70–72). Importantly, we ruled out that associations between our behavioural predictors and infant brain FC were driven by SES in our sample (see Supplementary Materials). Recent work further demonstrates that prenatal social disadvantage is associated with alterations in neonatal limbic connectivity that predict later emotional and behavioural outcomes (73). In this context, our findings suggest that interventions supporting caregiver–infant engagement and affectionate touch in the neonatal period could influence the developmental trajectories of functional networks implicated in social and affective processing, with potential downstream consequences for socioemotional development.

While the present findings highlight associations between caregiver behaviour and early functional connectivity, they cannot disentangle directionality. It remains an open question whether early social experience drives changes in brain organization, whether rapid changes in connectivity influence later interaction patterns, or whether these processes are bidirectional. Longitudinal studies extending beyond the neonatal period will be essential to determine the continuity of early caregiving-related differences in FC, as well as whether they predict later behavioural and cognitive outcomes.

## Materials and Methods

### 1.3 Participants

Participants were drawn from the Perinatal Imaging in Partnership with Families (PIPKIN) study, a UK-based longitudinal project examining how early life experiences and family environments influence infant development (www.pipkinstudy.com) (50). Ethical approval was granted by the Cambridge Psychology Research Ethics Committee (PRE.2020.123) and the East Midlands - Nottingham 1 Research Ethics Committee, on behalf of the NHS Health Research Authority (IRAS project ID: 295027, REC ref. 22/EM/0004). Parents provided written informed consent prior to participation. The present study drew on a subsample of the PIPKIN cohort who contributed data at the 1-month timepoint. For analyses of concurrent associations between FC and PCI, infants that underwent home sessions at 1 month (n = 80) were required to have both valid HD-DOT and PCI data. This resulted in a final analytic sample of n = 43 participants (25 male; 18 female). Of these, n = 37 (22 male; 15 female) participants have also provided valid FC data at least at another timepoint (either 1^st^ week, 2^nd^ week or both), thus contributing to the longitudinal analyses. Descriptives of the sample are provided in the SM (Supplementary Table 1 and Supplementary Figure 2) along with detailed reasons for exclusions for each data type (Supplementary Figure 1).

### 1.4 Data Acquisition

#### Functional connectivity

Task-free FC data were collected during home visits at one month using HD-DOT, while infants were sleeping and being held by an experimenter. The cap was positioned with both ears visible and the top edge above the eyebrows. We used the LUMO HD-DOT system (Gowerlabs Ltd, UK; https://www.gowerlabs.co.uk/lumo), comprising 12 tiles (each with 4 photodiode detectors and 3 dual-wavelength sources) covering frontal and temporal regions. Signal quality was monitored in real time, with a target of at least three motion-free 100-second segments (mean recording duration: 972 seconds, SD = 354). We aimed for ~10 minutes of clean data, as extended durations allow for the capture of low-frequency fluctuations, improve FC stability, sensitivity, and precision (74, 75).

#### Parent-infant interactions

Parent–infant interaction data were acquired during a 10-minute free-play session conducted in the home. The caregiver was instructed to interact with the infant as they normally would, without toys, while maintaining a seating position that kept both faces visible to the cameras while ensuring that parents felt at ease and comfortable. A particular strength of this study is that parent-infant interactions were observed in a more naturalistic setting than usual, indeed it was carried out at home rather than in a laboratory or hospital environment, allowing us to identify associations with behaviours as they occur in everyday interactions. The interaction was video-recorded simultaneously from three viewpoints using two high-definition Sony cameras and a webcam, positioned on individual tripods. One camera captured the caregiver’s face, the second captured the infant’s face, and the webcam provided a wide view of the overall scene. When recording started, the experimenter produced a hand clap within view of all cameras. This audiovisual event was used to synchronise the video streams during offline coding. The experimenter then left the room whenever possible to minimise interference with the interaction.

### 1.5 Data Processing

Data processing was performed in MATLAB using DOTHUB and Homer2 toolboxes, as well as Toast++ (76, 77). The pipeline entailed data preprocessing, image reconstruction and FC computation, and it is described below. Full function names and parameter settings are detailed in the Supplementary Materials.

#### 1.5.1 HD-DOT data

##### 1.5.1.1 Preprocessing

Preprocessing focused on identifying data segments ≥100 seconds with minimal motion artifacts. Motion artifacts were identified in short separation channels (<12mm). Channels where >90% of the recording was identified as motion were discarded. Time points were retained if >90% of short-separation channels were motion-free, and continuous segments ≥100 seconds were extracted. To maximize the duration of clean segments, and to include additional segments, brief motion artifacts (<2 sec) were corrected using wavelet denoising (78). Low-amplitude channels (dRange = [1e-4 2.5]) were also excluded. Within each segment, a 60-second window with maximal channel retention was automatically selected, using a signal-to-noise ratio of 12.5 (79). Intensity data were converted to optical density, then to HbO and HbR concentrations. Short-separation regression (SSR) was applied to attenuate the impact of extracerebral confounds (80). Bandpass filtering was performed using a linear regression model where sine and cosine functions modelled physiological signals (>0.08 Hz) (81), while Legendre polynomials accounted for fluctuations at very low frequencies (82). Data were converted back to optical density and down sampled to 1/10^th^ of its original frequency to optimize file size. Final datasets per participant were created by concatenating the normalised signal of each clean segment (concatenated duration: mean = 11.24 min, SD = 3.98), retaining only channels that were consistently preserved across all clean segments (channels included per wavelength: mean = 492.60, SD = 95.12).

Participants with <250 seconds of usable data were excluded, consistent with prior infant fNIRS FC studies (83–86). Participants could also be excluded due to insufficient channel retention (Supplementary Figure 9). Cap placement was evaluated from multi-angle session photos using a three-tier index relative to the forehead (Supplementary Figure 10), informing tilt correction during spatial registration. Full methods are presented in Supplementary Materials.

##### 1.5.1.2 Image Reconstruction

We selected a neonatal head model (37.9 cm, 44.28 weeks) from the Collins-Jones et al. (87) database to match our cohort’s (37.9 cm, 44.3 weeks). Optode and landmark coordinates were projected onto the model for individual-level reconstruction. Sensitivity profiles (Jacobians) were computed via finite element modeling in Toast++ (77). HbO/HbR concentration changes were reconstructed using zeroth-order Tikhonov regularization (hyperparameter = 0.01). For each participant, a binary sensitivity mask was created by thresholding the normalized Jacobians at 10% of the maximum. These were summed across participants, and a group-level mask was created by retaining nodes sensitive in ≥80% of participants (86). This mask was applied to all further analyses to constrain results to reliably sampled cortical regions. A cortical parcellation approach (detailed in the Supplementary Materials) was employed to aid seed selection.

##### 1.5.1.3 Functional Connectivity Computation

For each functional network, one left- and one right-hemisphere cortical seed were defined a priori and placed in the precentral gyrus (motor network), supramarginal gyrus (temporoparietal junction, TPJ), and superior frontal cortex (frontal network). For each seed, a surface-based region of interest was defined as all cortical mesh nodes within a 10-mm Euclidean radius. Euclidean distances from the seed to all cortical nodes were computed on the surface mesh and used to classify connections as short-(≤30 mm), medium-(30–60 mm), or long-range (60–120 mm), following Wang et al. (2021)(51).

For each participant, chromophore (HbO, HbR), network, and seed hemisphere, analyses were restricted to grey-matter–projected nodes surviving both individual- and group-level sensitivity masking. Seed signals were computed by averaging values across nodes within each seed region. Seed-to-whole-cortex functional connectivity (FC) was estimated using Pearson correlation between the seed signal and all remaining cortical nodes, yielding node-wise correlation coefficients (r), which were Fisher z-transformed for statistical analyses.

Group-level statistical masks were computed separately for each seed by performing node-wise one-sample t-tests on Fisher z-transformed FC values across participants, excluding seed nodes and nodes with <90% subject coverage, and applying false discovery rate (FDR) correction (q = 0.001). These masks were applied to individual FC maps. Then, weak connections were further excluded using subject-specific thresholding to account for inter-individual variability in overall FC magnitude. As choosing a threshold is arbitrary and might impact the results, we adopted a multiverse approach, which explicitly acknowledges this analytical flexibility and provides a sensitivity analysis that quantifies the impact of threshold-dependent processing decisions on statistical outcomes (88, 89). Specifically, metrics were recomputed across a set of percentile-based cut-offs (30–70%, in 10% increments) of the subject-specific positive FC distribution, with connections below threshold excluded.

Following group-level masking and individual-level thresholding, distance-based connectivity metrics were computed for each chromophore, network, and seed hemisphere. Connectivity strength was defined as the mean Fisher z-transformed FC within each distance category, and connectivity extent as the number of surviving connections within each category. The number of surviving connections by distance to each cortical parcel was also computed.

Two metrics were carried forward for analyses: connectivity strength and connectivity extent. For each participant, connectivity strength was defined as the Fisher z-transformed FC within each network, averaged across left and right hemispheres. Connectivity extent captured the total number of supra-threshold connections, computed as the sum of short-, medium-, and long-range connections and averaged across hemispheres. Connectivity within the seed was not included.

#### 1.5.2 Parent-infant interactions

Interactions were coded offline using Datavyu 1.3.8 (90) at 10 frames per second. Micro-analytic coding captured the onset and offset of gaze and touch behaviours, with coding labels adapted from existing coding schemes (Caregiver-Infant Touch Scale (91); Mother-Infant Touch Scale (92); Global Rating Scale (93); for review see (36)). For gaze, we separately coded instances in which one was looking at the other’s face, as well as looking away. For touch, we coded the onset and offset of affectionate tactile behaviours encompassing caress-like touch, hand holding, kissing, and playful touch. We then computed behavioural indices expressed as percentages of total interaction time: infant-to-parent gaze, parent-to-infant gaze, mutual gaze, affectionate touch incidence, and affectionate touch occurring while the infant was looking at the parent’s face. Inter-rater reliability was assessed on a randomly selected subset of participants (n = 13, 25% of sample) coded independently by two raters and was excellent (ICC = 0.979, 95% CI [0.968, 0.986]) (94).

For analyses, a dyadic engagement composite score was computed by summing infant-to-parent gaze, parent-to-infant gaze, and touch-to-boost incidences, capturing reciprocal engagement via both gaze and touch. Mutual gaze was not included in this index because it was primarily driven by infant behaviour and could bias the composite towards infant behaviour (Supplementary Figure 4, Supplementary Table 2). Affectionate touch incidence was analysed as a general index of tactile engagement, independent of infant looking behaviour. Additional detail on index computation, is provided in the Supplementary Materials.

### 1.6 Statistical Analysis

Statistical analyses were performed using R (R Core Team, 2013). For each dependent variable, linear mixed-effect models were built using the *lmer()* function from the lme4 package (95) and compared using *Akaike Information Criterion (AIC)* via the *compare_performance()* function (96). The model with the lowest AIC was considered the best-fitting, with AIC weights used to quantify relative model plausibility (97). We examined the best model’s fixed-effect estimates, p-values, and *R*^*2*^ values (marginal and conditional). Mixed models accounted for fixed effects (see Table 1 for list of variables) and included participant as a random effect to capture individual variability (98, 99). Full details of models comparisons including metrics of model performance and are provided in Supplementary Table 3. Model comparison sets were run separately for HbO and HbR.

**Table 1.**
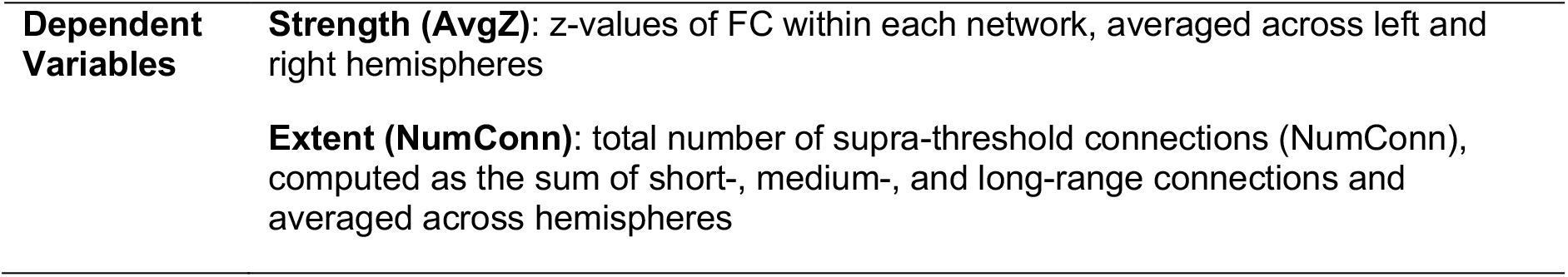
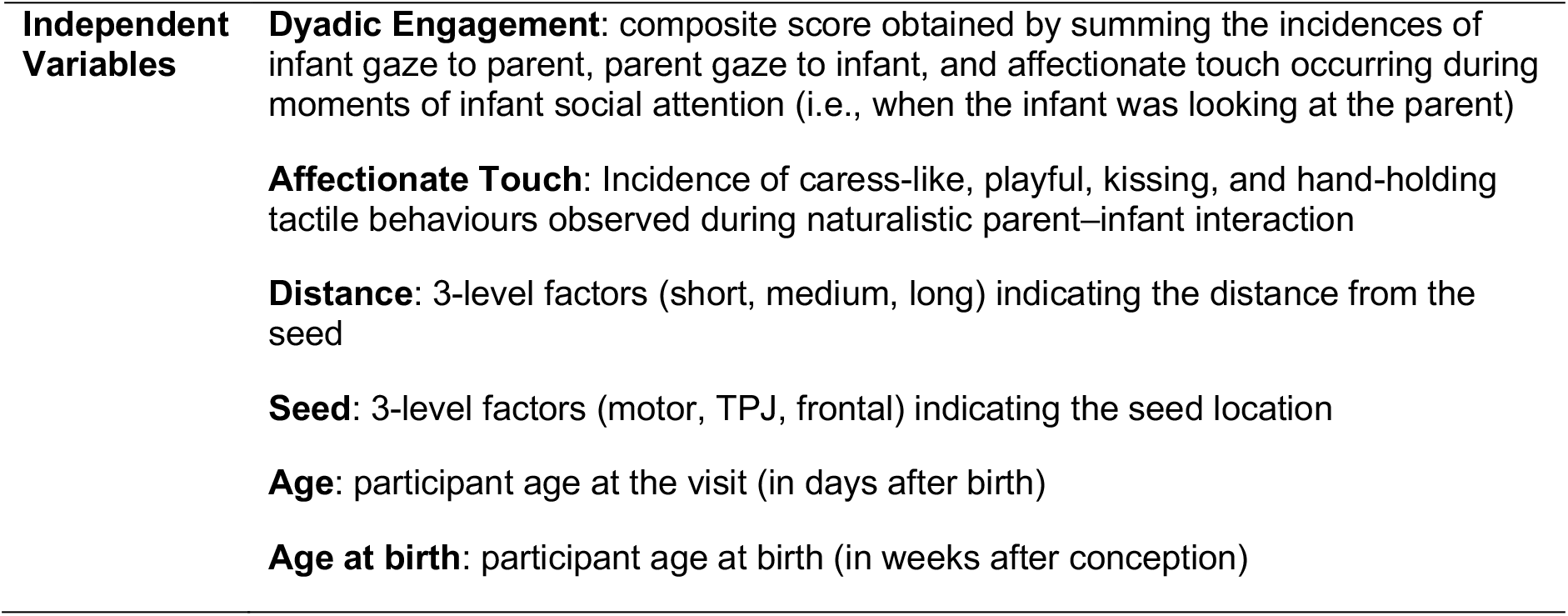
List of Dependent and Independent Variables.

| Dependent Variables | Strength (AvgZ): z-values of FC within each network, averaged across left and right hemispheres |
| --- | --- |
|  | Extent (NumConn): total number of supra-threshold connections (NumConn), computed as the sum of short-, medium-, and long-range connections and averaged across hemispheres |
| **Independent Variables** | **Dyadic Engagement:** composite score obtained by summing the incidences of infant gaze to parent, parent gaze to infant, and affectionate touch occurring during moments of infant social attention (i.e., when the infant was looking at the parent)<br>**Affectionate Touch:** Incidence of caress-like, playful, kissing, and hand-holding tactile behaviours observed during naturalistic parent–infant interaction<br>**Distance:** 3-level factors (short, medium, long) indicating the distance from the seed<br>**Seed:** 3-level factors (motor, TPJ, frontal) indicating the seed location<br>**Age:** participant age at the visit (in days after birth)<br>**Age at birth:** participant age at birth (in weeks after conception) |

For clarity of presentation, results are reported for HbO connectivity at a threshold excluding the weakest 30% of connections. The consistency of results across alternative thresholding scenarios and for HbR connectivity is reported in Supplementary Figure 18. Higher retention thresholds (e.g., excluding the weakest 30–40% of connections) preserve weaker and potentially long-range connections but may increase noise, whereas more stringent thresholds (e.g., excluding the weakest 50–70%) emphasize the strongest connections at the cost of number of connections retained and potential fragmentation biasing towards shorter distances (Supplementary Table 2; Supplementary Figure 13).

## Funding

BB and MJ were supported by a Medical Research Council Programme Grant (MR/T003057/1). LC and SLF were supported by a UKRI Future Leaders fellowship (MR/S018425/1).

## Acknowledgments

We are deeply grateful to all the families who generously gave their time to participate in this study. We would also like to thank Zayna Mian and Tridha Haritwal for their support with video coding.

## Author Contributions

**LC:** methodology, investigation, data curation, software, formal analysis, visualisation, writing – original paper. **BB:** methodology, investigation, data curation, software, formal analysis, writing – review and editing. **MR:** methodology, investigation, writing – review and editing. **MW:** investigation, writing – review and editing. **MJ:** conceptualisation, funding acquisition, supervision, writing – review and editing. **SLF:** conceptualisation, funding acquisition, methodology, project administration, supervision, writing – review and editing

## Competing Interest Statement

The authors declare that they have no competing interests.

## Data Availability Statement

The dataset is unique in that it provides the ability to link data points together (i.e. fNIRS data with outcome data or contextual factor data), but due to the nature of the data being collected (i.e. collected from a specific geographical location, longitudinal dataset of several datapoints) the majority of the data cannot be fully de-identified under the guidance included in the European General Data Protection Regulation (GDPR), and the conditions of our ethics approval do not allow public archiving of pseudonymised study data. Indeed, while fNIRS values after preprocessing and PCI summary values can be anonymised and will be made available in line with the PIPKIN protocol (Clackson et al., 2025), details of where the data is available can be found on the project OSF page: https://osf.io/tyh9q/overview. Some of the demographic information and videos used in this work cannot be deidentified for public archiving.

However, collaborations are encouraged, and projects are evaluated primarily on their consistency with the ethical principles and aims of the project that the families signed up to when partaking in this study. To access the data, interested readers should contact the PIPKIN coordinator on our website (https://www.pipkinstudy.com/people). Access will be granted to named individuals following ethical procedures governing the reuse of sensitive data. Requestors must complete and sign a data sharing agreement to ensure data is stored securely and used in accordance with the PIPKIN Project’s policies on Ethics, Data Sharing, Authorship and Publication.

## Notes

https://osf.io/tyh9q/overview

### Competing Interest Statement

The authors have declared no competing interest.

## References

1. M. H. Johnson, Interactive Specialization: A domain-general framework for human functional brain development? Developmental Cognitive Neuroscience 1, 7–21 (2011).

2. Karmiloff-Smith, M. S. Thomas, M. H. Johnson, Thinking developmentally from constructivism to neuroconstructivism. Milton Park: Routledge (2018).

3. D. Holland, et al., Structural Growth Trajectories and Rates of Change in the First 3 Months of Infant Brain Development. JAMA Neurol 71, 1266–1274 (2014).

4. S. Atzil, W. Gao, I. Fradkin, L. F. Barrett, Growing a social brain. Nat Hum Behav 2, 624– 636 (2018).

5. W. Gao, et al., Functional Network Development During the First Year: Relative Sequence and Socioeconomic Correlations. Cerebral Cortex 25, 2919–2928 (2015).

6. L. Carnevali, L. Della Longa, T. Farroni, M. H. Johnson, The neonatal brain at rest: A systematic review of task-free functional connectivity in the first 100 days. Neuroscience & Biobehavioral Reviews 186, 106703 (2026).

7. E. M. McCormick, J. van Hoorn, J. R. Cohen, E. H. Telzer, Functional connectivity in the social brain across childhood and adolescence. Soc Cogn Affect Neurosci 13, 819–830 (2018).

8. M. Fouladivanda, et al., Multi-scale structural rich-club organization of the brain in full-term newborns: a combined DWI and fMRI study. J Neural Eng 18 (2021).

9. P. Fransson, et al., Resting-state networks in the infant brain. PNAS Proceedings of the National Academy of Sciences of the United States of America 104, 15531–15536 (2007).

10. P. Fransson, U. Aden, M. Blennow, H. Lagercrantz, The functional architecture of the infant brain as revealed by resting-state fMRI. Cereb Cortex 21, 145–154 (2011).

11. M. F. Molloy, Z. M. Saygin, Individual variability in functional organization of the neonatal brain. NeuroImage 253, 119101 (2022).

12. O. Rajasilta, et al., Resting-state networks of the neonate brain identified using independent component analysis. Developmental Neurobiology 80, 111–125 (2020).

13. M. Sylvester, et al., Network-specific selectivity of functional connections in the neonatal brain. Cereb Cortex 33, 2200–2214 (2023).

14. J. Ciarrusta, et al., Emerging functional connectivity differences in newborn infants vulnerable to autism spectrum disorders. Transl Psychiatry 10, 131 (2020).

15. M. H. Johnson, A. Senju, P. Tomalski, The two-process theory of face processing: Modifications based on two decades of data from infants and adults. Neuroscience & Biobehavioral Reviews 50, 169–179 (2015).

16. S. R. Brindley, et al., Functional brain connectivity during social attention predicts individual differences in social skill. Social Cognitive and Affective Neuroscience 18, nsad055 (2023).

17. T. Farroni, et al., Infant cortex responds to other humans from shortly after birth. Sci. Rep. 3 (2013).

18. S. Lloyd-Fox, et al., Cortical specialisation to social stimuli from the first days to the second year of life: A rural Gambian cohort. Developmental Cognitive Neuroscience 25, 92–104 (2017).

19. H. Li, Z. Zhao, S. Jiang, H. Wu, Brain circuits that regulate social behavior. Mol Psychiatry 30, 3240–3256 (2025).

20. R. Feldman, Parent–infant synchrony: Biological foundations and developmental outcomes. Current directions in psychological science 16, 340–345 (2007).

21. I. Nomikou, K. J. Rohlfing, J. Szufnarowska, Educating attention: Recruiting, maintaining, and framing eye contact in early natural mother–infant interactions. Interaction Studies 14, 240–267 (2013).

22. M. Mercuri, et al., Mothers’ and fathers’ early tactile contact behaviors during triadic and dyadic parent-infant interactions immediately after birth and at 3-months postpartum: Implications for early care behaviors and intervention. Infant Behavior and Development 57, 101347 (2019).

23. G. M. Mason, M. H. Goldstein, J. A. Schwade, The role of multisensory development in early language learning. J Exp Child Psychol 183, 48–64 (2019).

24. Y. Okumura, Y. Kanakogi, T. Kobayashi, S. Itakura, Ostension affects infant learning more than attention. Cognition 195, 104082 (2020).

25. E. Parise, G. Csibra, Neural Responses to Multimodal Ostensive Signals in 5-Month-Old Infants. PLOS ONE 8, e72360 (2013).

26. T. Farroni, G. Csibra, F. Simion, M. H. Johnson, Eye contact detection in humans from birth. Proceedings of the National Academy of Sciences 99, 9602–9605 (2002).

27. T. Farroni, et al., Newborns’ preference for face-relevant stimuli: Effects of contrast polarity. Proceedings of the National Academy of Sciences 102, 17245–17250 (2005).

28. T. Farroni, S. Massaccesi, E. Menon, M. H. Johnson, Direct gaze modulates face recognition in young infants. Cognition 102, 396–404 (2007).

29. S. Rigato, E. Menon, V. D. Gangi, N. George, T. Farroni, The role of facial expressions in attention-orienting in adults and infants. International Journal of Behavioral Development 37, 154–159 (2013).

30. L. Carnevali, L. Della Longa, D. Dragovic, T. Farroni, Touch and look: The role of affective touch in promoting infants’ attention towards complex visual scenes. Infancy 29, 271–283 (2024).

31. L. Della Longa, T. Gliga, T. Farroni, Tune to touch: Affective touch enhances learning of face identity in 4-month-old infants. Developmental Cognitive Neuroscience 35, 42–46 (2019).

32. L. Della Longa, L. Carnevali, E. Patron, D. Dragovic, T. Farroni, Psychophysiological and Visual Behavioral Responses to Faces Associated with Affective and Non-affective Touch in Four-month-old Infants. Neuroscience (2020).

33. R. Feldman, A. I. Eidelman, L. Sirota, A. Weller, Comparison of Skin-to-Skin (Kangaroo) and Traditional Care: Parenting Outcomes and Preterm Infant Development. Pediatrics 110, 16–26 (2002).

34. Beebe, et al., A systems view of mother-infant face-to-face communication. Dev Psychol 52, 556–571 (2016).

35. S. Carozza, V. Leong, The Role of Affectionate Caregiver Touch in Early Neurodevelopment and Parent-Infant Interactional Synchrony. Front Neurosci 14, 613378 (2020).

36. Ilyka, M. H. Johnson, S. Lloyd-Fox, Infant social interactions and brain development: A systematic review. Neuroscience & Biobehavioral Reviews 130, 448–469 (2021).

37. K. Keunen, S. J. Counsell, M. J. N. L. Benders, The emergence of functional architecture during early brain development. NeuroImage 160, 2–14 (2017).

38. M. Seraji, et al., Spatial development of brain networks during the first six postnatal months. Commun Biol 8, 1514 (2025).

39. H. Zhang, D. Shen, W. Lin, Resting-state Functional MRI Studies on Infant Brains: a Decade of Gap-Filling Efforts. Neuroimage 185, 664–684 (2019).

40. J. Ciarrusta, R. Dimitrova, G. McAlonan, Early maturation of the social brain: How brain development provides a platform for the acquisition of social-cognitive competence. Prog Brain Res 254, 49–70 (2020).

41. M. H. Johnson, T. Grossmann, K. C. Kadosh, Mapping functional brain development: Building a social brain through interactive specialization. Developmental Psychology 45, 151–159 (2009).

42. T. Grossmann, M. H. Johnson, The development of the social brain in human infancy. European Journal of Neuroscience 25, 909–919 (2007).

43. M. H. Johnson, et al., The emergence of the social brain network: Evidence from typical and atypical development. Dev Psychopathol 17, 599–619 (2005).

44. W. Gao, W. Lin, K. Grewen, J. H. Gilmore, Functional Connectivity of the Infant Human Brain: Plastic and Modifiable. The Neuroscientist 23, 169 (2016).

45. R. Blanchett, et al., Genetic and environmental factors influencing neonatal resting-state functional connectivity. Cereb Cortex 33, 4829–4843 (2023).

46. Z. K. Mian, et al., Maternal Perinatal Depression, but not Anxiety, Associates with Infants’ Functional Connectivity at One Month of Age. [Preprint] (2025). Available at: https://osf.io/preprints/psyarxiv/3q6g4_v1/ [Accessed 7 November 2025].

47. C. M. Kelsey, K. Farris, T. Grossmann, Variability in Infants’ Functional Brain Network Connectivity Is Associated With Differences in Affect and Behavior. Frontiers in Psychiatry 12 (2021).

48. J. R. Chajes, J. A. Stern, C. M. Kelsey, T. Grossmann, Examining the Role of Socioeconomic Status and Maternal Sensitivity in Predicting Functional Brain Network Connectivity in 5-Month-Old Infants. Frontiers in Neuroscience (2022). 10.3389/FNINS.2022.892482.

49. J. Brauer, Y. Xiao, T. Poulain, A. D. Friederici, A. Schirmer, Frequency of Maternal Touch Predicts Resting Activity and Connectivity of the Developing Social Brain. Cereb Cortex 26, 3544–3552 (2016).

50. K. Clackson, et al., Perinatal Imaging in Partnership with Families (PIPKIN): Longitudinal cohort study protocol. PLOS ONE 20, e0330915 (2025).

51. Q. Wang, et al., Individual uniqueness in the neonatal functional connectome. Cerebral Cortex 31, 3701–3712 (2021).

52. T. Nguyen, D. H. Abney, D. Salamander, B. I. Bertenthal, S. Hoehl, Proximity and touch are associated with neural but not physiological synchrony in naturalistic mother-infant interactions. NeuroImage 244, 118599 (2021).

53. S. V. Wass, M. Whitehorn, I. Marriott Haresign, E. Phillips, V. Leong, Interpersonal Neural Entrainment during Early Social Interaction. Trends in Cognitive Sciences 24, 329–342 (2020).

54. S. Grayson, D. A. Fair, Development of large-scale functional networks from birth to adulthood: a guide to neuroimaging literature. Neuroimage 160, 15–31 (2017).

55. Shi, A. P. Salzwedel, W. Lin, J. H. Gilmore, W. Gao, Functional Brain Parcellations of the Infant Brain and the Associated Developmental Trends. Cereb Cortex 28, 1358–1368 (2018).

56. A. N. Nielsen, et al., Maturation of large-scale brain systems over the first month of life. Cereb Cortex 33, 2788–2803 (2023).

57. Vahidi, et al., Investigating Task-Free Functional Connectivity Patterns in Newborns Using functional Near-Infrared Spectroscopy. [Preprint] (2023). Available at: https://www.biorxiv.org/content/10.1101/2023.09.05.555980v1 [Accessed 21 September 2024].

58. M. Cao, H. Huang, Y. He, Developmental Connectomics from Infancy through Early Childhood. Trends Neurosci 40, 494–506 (2017).

59. T. Grossmann, Novel Insights into the Social Functions of the Medial Prefrontal Cortex during Infancy. eNeuro 12 (2025).

60. T. Grossmann, The social self in the developing brain. Neuroscience & Biobehavioral Reviews 169, 106023 (2025).

61. M. S. Scher, et al., Neurophysiologic assessment of brain maturation after an 8-week trial of skin-to-skin contact on preterm infants. Clin Neurophysiol 120, 1812–1818 (2009).

62. C. Schneider, N. Charpak, J.G. Ruiz-Peláez, R. Tessier, Cerebral motor function in very premature-at-birth adolescents: a brain stimulation exploration of kangaroo mother care effects. Acta Paediatr 101, 1045–1053 (2012).

63. V. Mateus, A. Osório, H. O. Miguel, S. Cruz, A. Sampaio, Maternal sensitivity and infant neural response to touch: an fNIRS study. Social cognitive and affective neuroscience (2021). 10.1093/SCAN/NSAB069.

64. W. Gao, A hierarchical model of early brain functional network development. Trends Cogn Sci 29, 855–868 (2025).

65. M. Imai, et al., Functional connectivity of the cortex of term and preterm infants and infants with Down’s syndrome. NeuroImage 85, 272–278 (2014).

66. M. Ouyang, H. Kang, J. A. Detre, T. P. L. Roberts, H. Huang, Short-range connections in the developmental connectome during typical and atypical brain maturation. Neuroscience & Biobehavioral Reviews 83, 109–122 (2017).

67. S. Dall’Orso, et al., Development of functional organization within the sensorimotor network across the perinatal period. Human Brain Mapping 43, 2249–2261 (2022).

68. M. Eyre, et al., The Developing Human Connectome Project: typical and disrupted perinatal functional connectivity. Brain 144, 2199–2213 (2021).

69. E. M. Gordon, et al., A somato-cognitive action network alternates with effector regions in motor cortex. Nature 617, 351–359 (2023).

70. P. Tomalski, M. H. Johnson, The effects of early adversity on the adult and developing brain. Current Opinion in Psychiatry 23, 233 (2010).

71. S. Nolvi, E. C. Merz, E.-L. Kataja, C. E. Parsons, Prenatal Stress and the Developing Brain: Postnatal Environments Promoting Resilience. Biological Psychiatry 93, 942–952 (2023).

72. J. L. Luby, et al., Basic Environmental Supports for Positive Brain and Cognitive Development in the First Year of Life. JAMA Pediatrics 178, 465–472 (2024).

73. M. P. Herzberg, et al., Maternal prenatal social disadvantage and neonatal functional connectivity: Associations with psychopathology symptoms at age 12 months. Developmental Psychology 60, 1562–1579 (2024).

74. K. R. A. Van Dijk, et al., Intrinsic functional connectivity as a tool for human connectomics: theory, properties, and optimization. J Neurophysiol 103, 297–321 (2010).

75. I. Paranawithana, D. Mao, C. M. McKay, Y. T. Wong, Connections between spatially distant primary language regions strengthen with age during infancy, as revealed by resting-state fNIRS. J Neural Eng 20 (2023).

76. T. J. Huppert, S. G. Diamond, M. A. Franceschini, D. A. Boas, HomER: a review of time-series analysis methods for near-infrared spectroscopy of the brain. Appl. Opt., AO 48, D280–D298 (2009).

77. M. Schweiger, S. R. Arridge, The Toast++ software suite for forward and inverse modeling in optical tomography. JBO 19, 040801 (2014).

78. A. X. Patel, et al., A wavelet method for modeling and despiking motion artifacts from resting-state fMRI time series. NeuroImage 95, 287–304 (2014).

79. E. M. Frijia, et al., Functional imaging of the developing brain with wearable high-density diffuse optical tomography: A new benchmark for infant neuroimaging outside the scanner environment. Neuroimage 225, 117490 (2021).

80. E. E. Vidal-Rosas, et al., Evaluating a new generation of wearable high-density diffuse optical tomography technology via retinotopic mapping of the adult visual cortex. Neurophotonics 8, 025002 (2021).

81. C. Caballero-Gaudes, R. C. Reynolds, Methods for cleaning the BOLD fMRI signal. NeuroImage 154, 128–149 (2017).

82. B. Blanco, et al., Group-level cortical functional connectivity patterns using fNIRS: assessing the effect of bilingualism in young infants. Neurophotonics 8, 025011 (2021).

83. B. Blanco, M. Molnar, M. Carreiras, C. Caballero-Gaudes, Open access dataset of task-free hemodynamic activity in 4-month-old infants during sleep using fNIRS. Sci Data 9, 102 (2022).

84. C. Bulgarelli, et al., The developmental trajectory of fronto-temporoparietal connectivity as a proxy of the default mode network: a longitudinal fNIRS investigation. Human brain mapping (2020). 10.1002/HBM.24974.

85. C. W. Lee, et al., Sleep State Modulates Resting-State Functional Connectivity in Neonates. Front. Neurosci. 14 (2020).

86. J. Uchitel, et al., Cot-side imaging of functional connectivity in the developing brain during sleep using wearable high-density diffuse optical tomography. NeuroImage 265, 119784 (2023).

87. L. H. Collins-Jones, et al., Construction and validation of a database of head models for functional imaging of the neonatal brain. Human Brain Mapping 42, 567–586 (2021).

88. Calignano, P. Girardi, G. Altoè, First steps into the pupillometry multiverse of developmental science. Behav Res 56, 3346–3365 (2024).

89. C. Montuori, F. Gambarota, G. Altoé, B. Arfé, The cognitive effects of computational thinking: A systematic review and meta-analytic study. Computers & Education 210, 104961 (2024).

90. Datavyu Team, Datavyu: A video coding tool. (2014). Deposited 2014.

91. D. M. Stack, A. D. L. Jean, “Communicating through touch: Touching during parent-infant interactions” in The Handbook of Touch: Neuroscience, Behavioral, and Health Perspectives, (Springer Publishing Company, 2011), pp. 273–298.

92. L. Crucianelli, et al., The mindedness of maternal touch: An investigation of maternal mind-mindedness and mother-infant touch interactions. Developmental Cognitive Neuroscience 35, 47–56 (2019).

93. L. Murray, A. Fiori-Cowley, R. Hooper, P. Cooper, The impact of postnatal depression and associated adversity on early mother-infant interactions and later infant outcome. Child Dev 67, 2512–2526 (1996).

94. K. A. Hallgren, Computing Inter-Rater Reliability for Observational Data: An Overview and Tutorial. Tutor Quant Methods Psychol 8, 23–34 (2012).

95. D. Bates, M. Mächler, B. Bolker, S. Walker, Fitting Linear Mixed-Effects Models Using lme4. Journal of Statistical Software 67, 1–48 (2015).

96. D. Lüdecke, M. S. Ben-Shachar, I. Patil, P. Waggoner, D. Makowski, performance: An R package for assessment, comparison and testing of statistical models. Journal of Open Source Software 6, 3139 (2021).

97. E.-J. Wagenmakers, S. Farrell, AIC model selection using Akaike weights. Psychonomic Bulletin & Review 11, 192–196 (2004).

98. R. H. Baayen, D. J. Davidson, D. M. Bates, Mixed-effects modeling with crossed random effects for subjects and items. Journal of Memory and Language 59, 390–412 (2008).

99. A. Gelman, J. Hill, Data Analysis Using Regression and Multilevel/Hierarchical Models. Higher Education from Cambridge University Press (2006). Available at: https://www.cambridge.org/highereducation/books/data-analysis-using-regression-and-multilevel-hierarchical-models/32A29531C7FD730C3A68951A17C9D983 [Accessed 21 September 2024].

