## Supplementary Materials for "Caregiver-infant interactions selectively shape emerging functional connectivity in the neonatal brain"

Table of Contents

***1. Participants*** ***2***
***2. PCI*** ***4***
**a. Data acquisition** **4**
**b. Data processing** **4**
***3. HD-DOT*** ***6***
**a. Data acquisition** **6**
**b. Detailed HD-DOT data processing** **6**
i. Preprocessing of HD-DOT data 6
ii. Image Reconstruction 9
iii. Cortical Parcellation 9
**c. Details on exclusion of neuroimaging datasets** **10**
**d. Cap positioning and three-tiered index for spatial registration** **10**
**e. Seed-based analyses and multiverse thresholding** **11**
***4. STATISTICAL ANALYSES*** ***16***
**a. Model comparisons** **16**
**b. Consistency of results across thresholding scenarios and chromophores** **18**
**c. Variance partitioning analyses to check influences on dyadic engagement composite** **19**
**d. Ruling out socioeconomic status as a confound** **20**

### 1. Participants

Eighty **1-month home sessions** were carried out as part of the PIPKIN study. Within these sessions, n = 52 infants provided codable infant-caregiver interaction data, with an additional n = 14 datasets collected but deemed unusable (technical issues: n = 4; infant fussiness: n = 5; infant asleep: n = 3; other reasons: n = 2). Valid task-free functional connectivity (FC) data were obtained from n = 60 participants, with an additional n = 19 datasets excluded during preprocessing (cap displacement: n = 4; inadequate recording duration: n = 13; insufficient number of channels: n = 2). A flowchart of number of participants with different reasons for exclusions across datatypes is shown in Supplementary Figure 1. For the final sample, a visualisation of descriptives for age, head circumference, FC clean recording duration and number of included channels per wavelength is in **Error! Reference source not found.**.

*Supplementary Figure 1.* Details on participants exclusion for each data type.

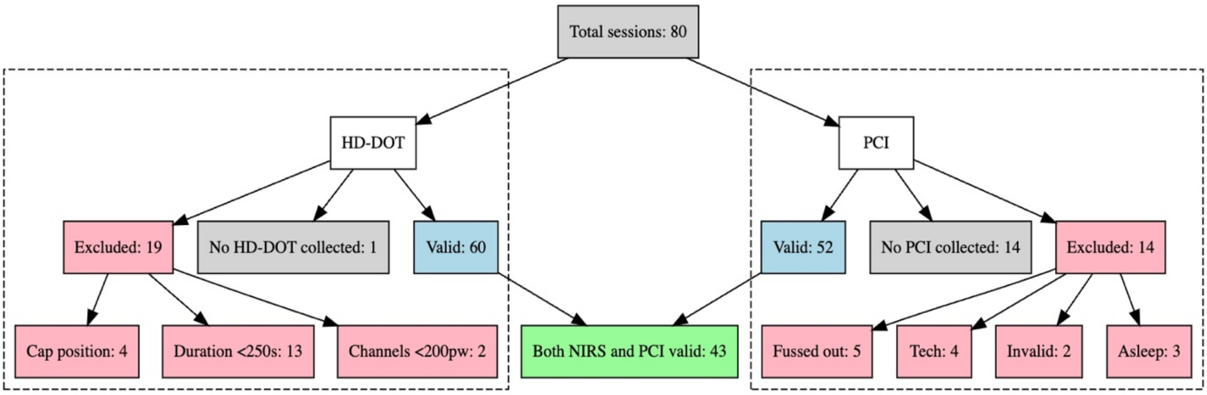


Therefore, n = 43 participants have valid FC and valid PCI at 1 month, contributing to the 1-month analyses. Of these, n = 37 participants have also provided valid FC data at least at another timepoint (either 1st week, 2nd week or both), thus contributing to the longitudinal analyses. Age descriptives are in Supplementary Table 1 and a visual of its distributions is provided in

Supplementary Figure 2

*Supplementary Table 1.* Age descriptives for the PIPKIN cohort sample in this study

| **Age at birth** | | **Age at test** | | | | |
|---|---|---|---|---|---|---|
| **Weeks post-conception** | | **Weeks post-conception** | | | **Days after birth** | |
| Mean (SD) | Range | | Mean (SD) | Range | Mean (SD) | Range |
| 1 month sample (n = 43) | | | | | | |
| 39.76 (1.12) | 37.29–41.86 | | 45.13 (1.58) | 41.86–49.14 | 37.6 (6) | 30–55 |
| Longitudinal sample (n = 37) | | | | | | |
| 39.82 (1.13) | 37.29–41.86 | 1st week | 40.93 (1.15) | 38.43–43.14 | 8.04 (2.55) | 4–14 |
| | | 2nd week | 42.66 (1.2) | 40.43–45 | 19.39 (4.81) | 9–31 |
| | | 1 month | 45.26 (1.57) | 42.57–49.14 | 38.05 (6.2) | 30–55 |

*Supplementary Figure 2.* Distribution of participants age at test across timepoints

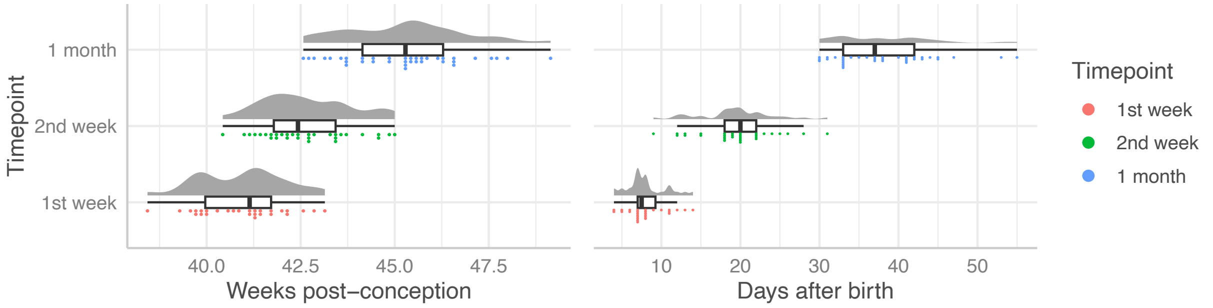


### 2. PCI

#### a. Data acquisition

Three cameras captured the interaction from complementary angles: a Sony camera on a short tripod focused on the caregiver's face; a second Sony camera on a tall tripod focused on the infant's face; and a fixed webcam mounted on a screen-height tripod providing an overview of the dyad and room. Prior to recording, the experimenter ensured that both faces were clearly visible with sufficient space for movement. Once all cameras were positioned, recording was initiated and parents received the following standardised instruction: "*Play with your child for around 10 minutes as you normally do, without toys. Please try to remain in this position so that the cameras retain a good view, unless your child becomes upset, in which case do whatever is needed.*" At the beginning of each recording, the experimenter clapped once while clearly visible to all cameras to facilitate audiovisual synchronisation during later offline coding. When possible, the experimenter left the room for the duration of the interaction; otherwise, they remained quietly out of view. Recordings were stopped after 10 minutes or earlier if needed (i.e. baby fussed out or fell asleep).

As part of the broader PIPKIN protocol, interaction was recorded between the infant and their mother (n = 66 datasets collected) as well as father (n = 29 datasets collected) where possible. In the present study, only data from mother-infant interaction were used.

#### b. Data processing

Interactions were coded in Datavyu 1.3.8 software, a video coding and data visualization tool for collecting behavioural data from video (Datavyu Team, 2014) at 10 frames per second. A micro-analytic coding scheme was used, whereby for each behaviour observed its onset and offset were recorded, for all their occurrences, to derive measures of behavioural duration. The coding scheme used to quantify the prevalence of specific gaze and touch behaviours in parent-infant dyads was adapted from existing coding schemes (Caregiver-Infant Touch Scale [1]; Mother-Infant Touch Scale [2] ; Global Rating Scale [3]; for review see [4]). For gaze behaviour, the amount of time the parent and infant each spent looking at the other's face (face looking) and away from the other's face (cut gaze) was coded. For the tactile domain, hand holding, kissing, playful and caress-like touch were coded as affectionate touch. Because the duration of recordings varied across dyads (in minutes: Mean = 5.83; SD = 1.46; Range = 2.30 - 8.55; median = 6.03; IQR = 4.92-6.45), all behavioural variables were normalized by the total interaction length and expressed as percentages of total interaction time.

Offline coding of a randomly selected subgroup of participants (n = 13, 25% of the total sample) was performed by two independent coders. Inter-rater reliability, calculated in R using the *icc()* function from *irr()* package [5], was found to be excellent (ICC = 0.979 [0.968; 0.986]) [6].

After behavioural coding, the data were exported to R to compute behavioural indexes of interest (see distributions in *Supplementary Figure 3*):

- Infant to parent = percentage of time when infant is looking at parent;
- Parent to infant = percentage of time when parent is looking at infant;
- Mutual gaze = percentage of time when parent and infant are looking at each other;
- Affectionate touch incidence = percentage of time when affectionate touch was occurring;

- Affectionate touch to boost = percentage of time when affectionate touch was occurring whilst the infant is looking at parent's face;
- Affectionate touch to engage = percentage of time when affectionate touch was occurring whilst the infant was not looking at parent's face.

*Supplementary Figure 3.* Distribution of percentage values for the various behaviours coded; respectively incidence of: mutual gaze, infant looking at parent's face, parent looking at infants' face, total affectionate touch, affectionate touch provided when infant was looking at parent's face.

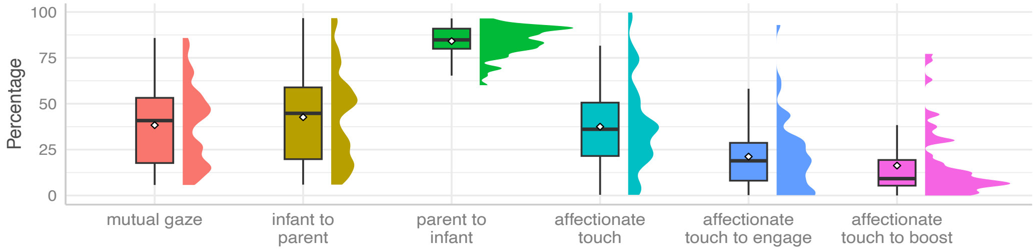


For the analyses, a **dyadic engagement** composite score capturing a dimension of reciprocal social engagement via both gaze and touch was computed by summing the normalised percentages of infant-to-parent gaze, parent-to-infant gaze, and affectionate touch occurring when infant was looking at parent. **Affectionate touch** was considered separately as a broader measure of tactile engagement, reflecting the percentage of total interaction time during which affectionate touch occurred regardless of the infant's attentional focus.

Whilst the creation of the above variables (dyadic engagement, affectionate touch) carried on for analyses was theoretically motivated, we also run a principal component analysis (PCA) on the above behavioural variables (mutual gaze, infant to parent, parent to infant, affectionate touch, affectionate touch to engage, affectionate touch to boost) to support this choice. The first two components explained 82.6 % of the total variance (PC1 = 49.9%, PC2 = 32.7%). Loadings of different behaviours on each composite are reported in Supplementary Table 2.

*Supplementary Table 2.* PCA loadings on parent-infant interaction behaviours. In bold loadings of behaviours that contributed to the scores used for further analyses

| **Behaviour** | **PCA Component** | |
|---|---|---|
| | **PC1** | **PC2** |
| Mutual gaze | 0.56 | -0.01 |
| Infant to parent gaze | **0.56** | -0.04 |
| Parent to infant gaze | **0.25** | 0.26 |
| Affectionate touch incidence | 0.05 | **-0.70** |
| Affectionate touch to boost (when infant engaged) | **0.43** | -0.42 |
| Affectionate touch when infant disengaged | -0.36 | -0.51 |

The first component (PC1) was characterised by strong loadings for mutual gaze (0.56) and infant-to-parent gaze (0.56), with additional contributions from affectionate touch occurring while the infant was attending to the parent (0.43) and parent-to-infant gaze (0.25) – indeed reflecting a dimension of reciprocal dyadic engagement. The comparable loadings of mutual gaze and infant-to-parent gaze indicate that variation in mutual gaze was largely driven by infant looking behaviour. For this reason, mutual gaze was not included in the dyadic engagement composite score, as doing so would have disproportionately weighted infant attention within the index. Notably, although variability in parent gaze was smaller relative to infant gaze and therefore the loading on the PC1 was lower, it was nevertheless present and meaningful. In fact, the positive loadings of parent-to-infant gaze and affectionate touch occurring during infant attention indicate that parental behaviours contribute additional variance to this interactional dimension. Therefore, the dyadic engagement composite combined infant-to-parent gaze, parent-to-infant gaze, and affectionate touch occurring while the infant was attending to the parent.

The second component (PC2) loaded most strongly on overall affectionate touch incidence (-0.70) and affectionate touch when the infant was disengaged (-0.51), with an additional contribution from affectionate touch when the infant was engaged (-0.42). Together, these loadings indicate a touch-focused dimension reflecting the general frequency of tactile engagement, independent of the infant's attentional state. This pattern justified treating overall affectionate touch incidence as a separate composite metric in subsequent analyses.

### 3. HD-DOT

#### a. Data acquisition

*Supplementary Figure 4.* Neuroimaging data acquisition with HD-DOT system (LUMO), with tiles (A), light guides (B) and headgear (C)

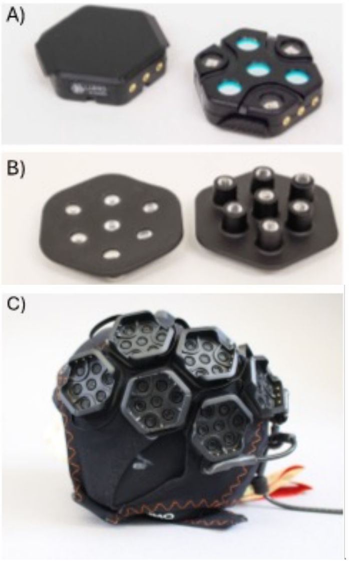


#### b. Detailed HD-DOT data processing

Data processing was performed in MATLAB using DOTHUB and Homer2 toolboxes, as well as Toast++ [7,8]. The pipeline entailed data preprocessing, image reconstruction, cortical parcellation and FC computation. Each of these steps is described below.

##### i. Preprocessing of HD-DOT data

A schematic representation of the preprocessing steps is provided in Supplementary Figure 5. The first step involved identifying and extracting clean data segments that were at least 100 seconds long. In an initial assessment of the data, motion artifact detection was performed on short separation channels (<12mm) using the *hmrMotionArtifactByChannel* function with the following parameters: tMotion = 0.5, tMask = 5, STDEVthresh = 10, AMPthresh = 0.1. Parameters were selected based on Uchitel et al. [9]. Those channels where more than 90% of the recording was identified as motion were rejected. Then, for each time sample we computed the percentage of motion-free channels, to identify time points where more than 90% of short-distance channels were motion-free. From this, clean segments that were at least 100 seconds long were extracted. To maximize the duration of clean segments, and to include additional segments, brief motion artifacts (less than 2 sec) were included and later corrected by means of wavelet denoising (Patel et al, 2014). Before this step, channels displaying low amplitude were excluded using *enPruneChannels* function (dRange = [1e-4 2.5]).

For each clean segment, channel rejection was performed with a customization of *enPruneChannels* function which uses a 60-second sliding window to automatically select a segment with the highest channel retention based on its signal-to-noise ratio of 12.5 (Frijia et al., 2021). A visualisation of channel retention across timepoints and source-detectors distances is provided in Supplementary Figure 6. Intensity data were then converted to optical density and oxy- and deoxyhaemoglobin (HbO and HbR) concentration changes using *hmrOD2Conc* function. Short-separation regression (SSR) was included to attenuate the impact of extracerebral confounds (*DOTHUBhmrSSRegressionByChannel*, flagSSmethod = 2, rhoSDssThresh = 10, [10]). This was followed by bandpass filtering, using a linear regression model to remove very low frequency fluctuations [11]. Sine and cosine functions for frequencies above 0.08 Hz were used to remove the contribution of physiological noise sources (e.g. respiration and cardiac pulsation), and Legendre polynomials were included to account for fluctuations at very low frequencies [12]. After filtering, we converted concentration data back to optical density for image reconstruction using *hmrConc2OD* function and down sampled to 1/10th of its frequency to optimize file size, allowing for separate files to be generated for each clean data segment. At this point, we used the *DOTHUBwritePREPRO* function to save a separate file for each preprocessed data segment. After preprocessing all the clean segments, their normalized signal was concatenated for those channels that were consistently retained across data segments, generating a single dataset for each participant.

Supplementary Figure 5. Schematic representation of preprocessing steps

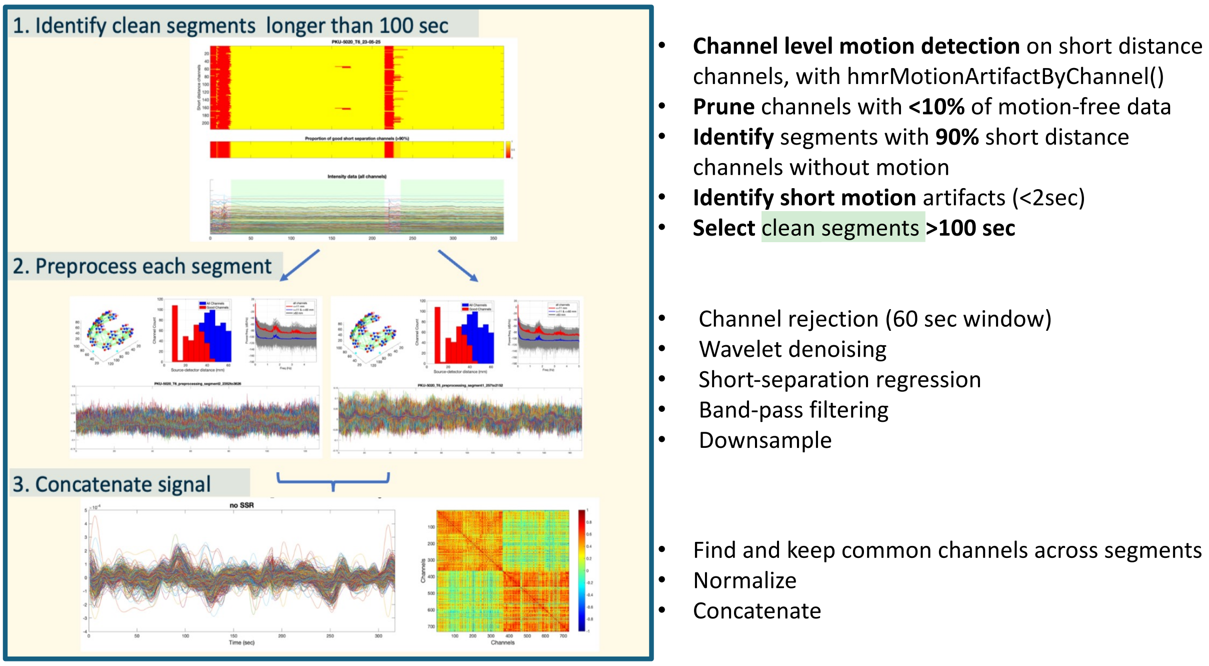

*Supplementary Figure 6.* For each of the timepoint examined, **A)** shows average counts of included channels (red) at various source-detector distances out of the total channels (blue); **B)** visualises such channels on our cap layout, with different plots for three groups of source-detector distance channels and graded yellow to green colour to represent the proportion of participants for which a certain channel was included.

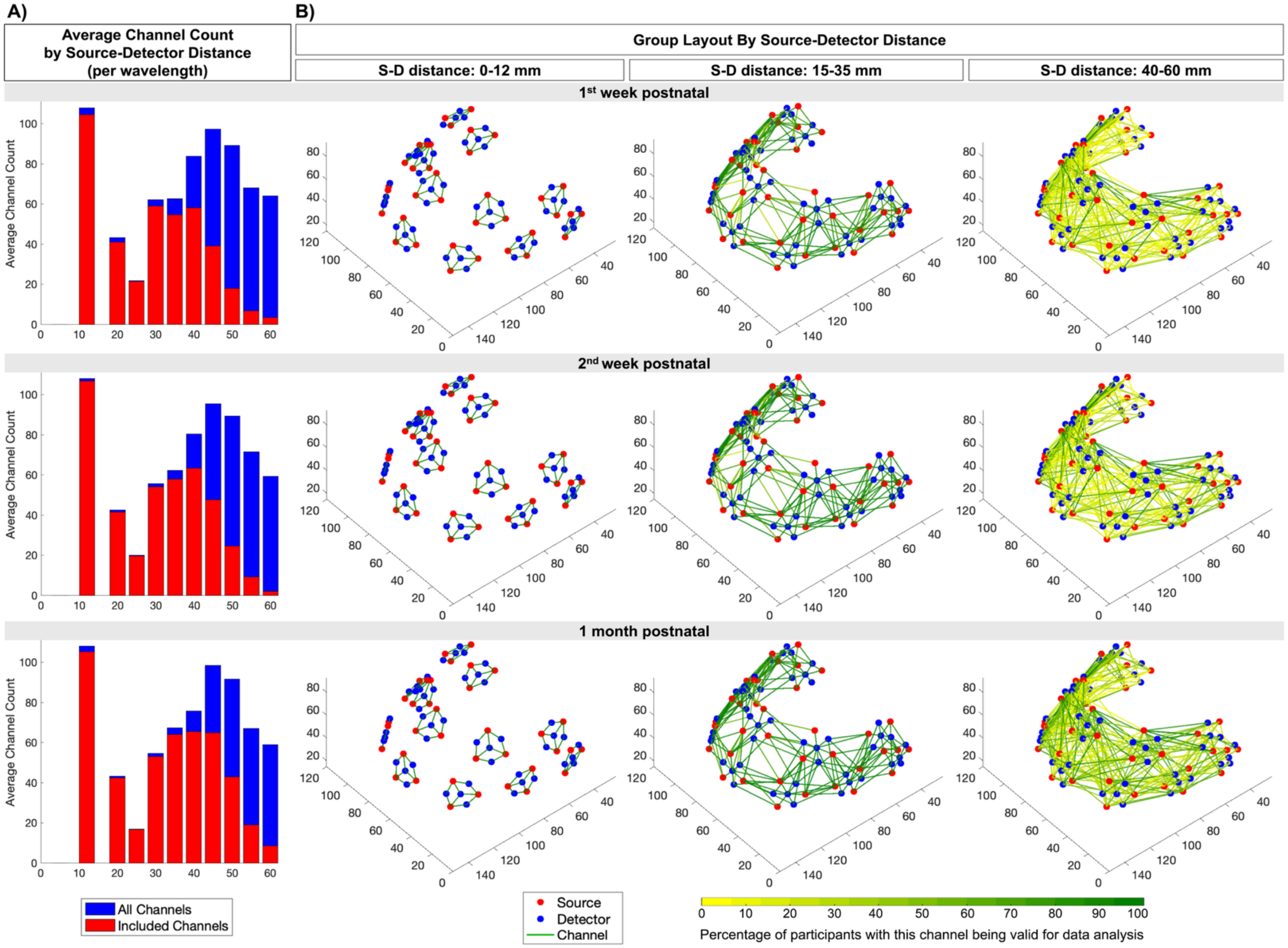

#### ii. Image Reconstruction

A single head model was used for image reconstruction across all participants. This model was selected from the database provided by Collins-Jones et al. [13], which includes a set of structural neonatal head models varying in head circumference and gestational age. To ensure anatomical relevance, we selected the head model whose combination of head circumference and gestational age most closely matched the average values observed in our sample (M = 37.88 cm, SD = 1.09, range: 35-39.7 cm). Specifically, the selected head model's head circumference measured 37.9 cm, with a gestational age of 44.28 weeks. Individual-specific optode and landmark coordinates, as estimated by our optode registration method, were projected onto the head model and used for subject-level image reconstruction. The sensitivity profile (Jacobian matrix) for each wavelength was computed using the finite element method in Toast++ [8]. A zeroth-order Tikhonov regularised reconstruction was performed using a regularisation hyperparameter of 0.01. Reconstructed images of changes in HbO and HbR concentrations were derived at each time point for every node of the grey matter mesh. The code employed for preprocessing and reconstructing the HD-DOT data is available as part of the DOT-HUB toolbox at www.github.com/DOT-HUB . A group-level sensitivity mask was calculated by finding the sensitivity for each surface node at the participant level using the *DOTHUB_MakeGMSensitivityMap* function. A node was considered sensitive if it exhibited a sensitivity greater than 10% the maximum value of the normalized Jacobian. For each participant, a binary sensitivity mask was computed, which was then summed across participants, and any nodes which were sensitive in >80% of participants were included in the final group-level mask [9].

*Supplementary Figure 7.* Group sensitivity mask, with colours representing the percentage of participants with that part of surface being covered

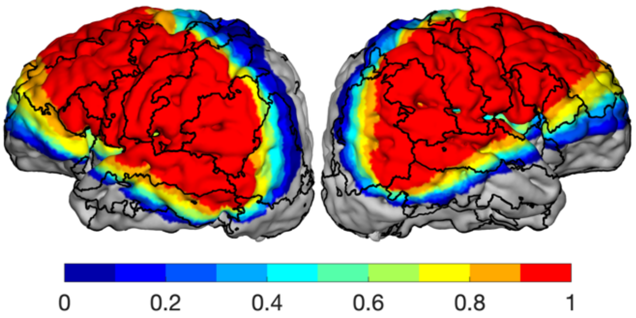


#### iii. Cortical Parcellation

A cortical parcellation approach was employed to aid seed selection. The M-CRIB neonatal atlas [14], including 10 neonatal T2-weighted MRI scans, was processed using the developing Human Connectome Project [15] pipeline to create brain masks and segment different tissue types. Then each atlas brain was aligned with the head model by matching the intensity distributions with the T2-weighted scans, after which each M-CRIB T2 volume was non-linearly registered to the head model using the ANTs toolbox [16]. Following this, the labels from all 10 atlas subjects were combined using a joint label fusion approach, resulting in a cortical parcellation with 70 regions of interest. The group-level sensitivity mask was applied to the cortical parcellation, both visualised in Supplementary Figure 8.

*Supplementary Figure 8*. Cortical parcellation masked by group sensitivity for the full FC sample (n = 60).

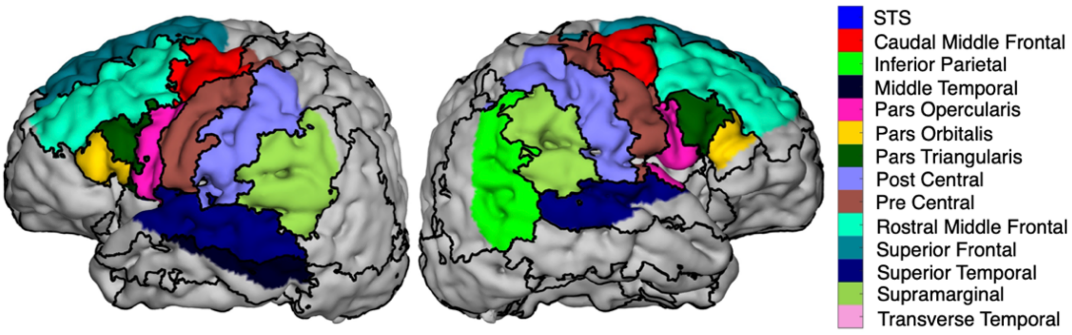

#### c. Details on exclusion of neuroimaging datasets

*Supplementary Figure 9.* (A) Distribution of clean data duration (seconds) among participants with FC datasets available at different timepoints and PCI run. Dotted line indicates the 250 sec threshold for inclusion. The one datapoint that is far from the others indicates one of the early babies tested, where we were working out for how long recording could go on, therefore for that participant we recorded for as much as they were happy to. (B) Distribution of retained channels per wavelength. Dotted line indicates the 200 channels threshold for inclusion.

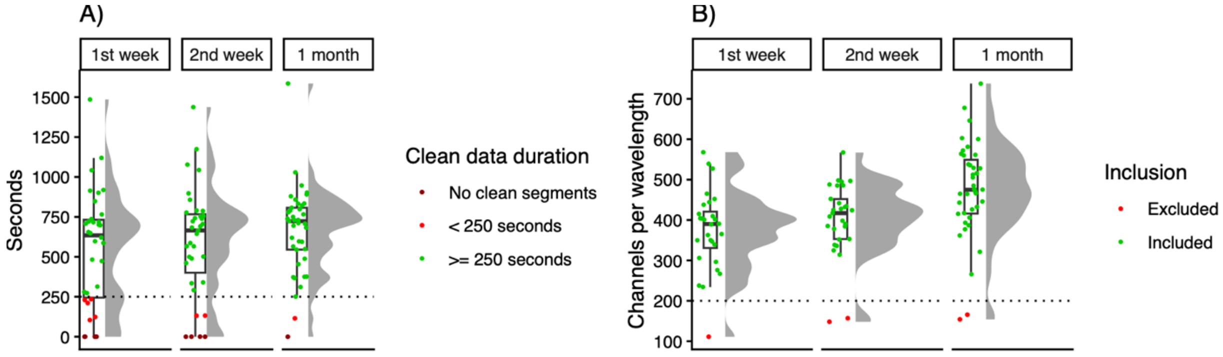


#### d. Cap positioning and three-tiered index for spatial registration

The quality of cap positioning was noted based on pictures acquired during the session from different angles and used to exclude participants when the cap was displaced. Cap placement was considered correct when placed above infants' eyebrows, with the tiles centred, ensuring coverage of frontal brain regions. HD-DOT datasets of participants with valid interaction data at 1 month could be excluded from analysis in case of poor cap placement or tiles not being centred (n=1 at 1$^{st}$ week, n = 4 at 2$^{nd}$ week, n=1 at 1 month). For participants with adequate cap positioning, a three-tiered index was used to classify the extent to which cap covered participant's forehead. Tier 1: Cap placed just above infants' eyebrows (ideal placement, n = 23). Tier 2: Cap placed in mid-frontal position (half of forehead visible, n = 19). Tier 3: Cap placed close to the hairline (more than half forehead visible, n = 9).

The three-tiered index was later used for registering source-detector positions to the head model. A custom MATLAB function was developed to compensate for variations in HD-DOT cap placement across participants. This function applies a smooth, graded backward rotation to the headband layout by rotating optode positions around a common posterior anchor point and a fixed axis (the x-axis). The degree of rotation for each tile in the layout is proportional to its anterior-posterior position, with the most anterior tiles rotated by the maximum specified angle and posterior tiles remaining nearly fixed. This approach preserves the spatial geometry of the optode layout and maintains consistent within tile structure, ensuring that the headband configuration accurately reflects differences in headband placement while minimising distortion of the original layout. Similar methods have been used in previous studies using HD-DOT with infant participants [17,18], for which accurate and easy to use method for spatial registration of optode locations are not yet available. Based on the cap positioning categories, fixed rotations of 7.5 degrees (Tier 2) and 15 degrees (Tier 3) were applied to approximate cap tilt.

As a sanity check, we made sure data quality was not impacted by cap position through visualising if the total clean duration and number of channels retained varied across the three tiers (Supplementary Figure 10).

*Supplementary Figure 10.* Metrics (A) Clean duration, (B) Number of retained channels per wavelength) by tier (Tier 1: neutral/original placement; Tier 2 = tilted by 7.5 degrees; Tier 3 = tilted by 15 degrees) across timepoints.

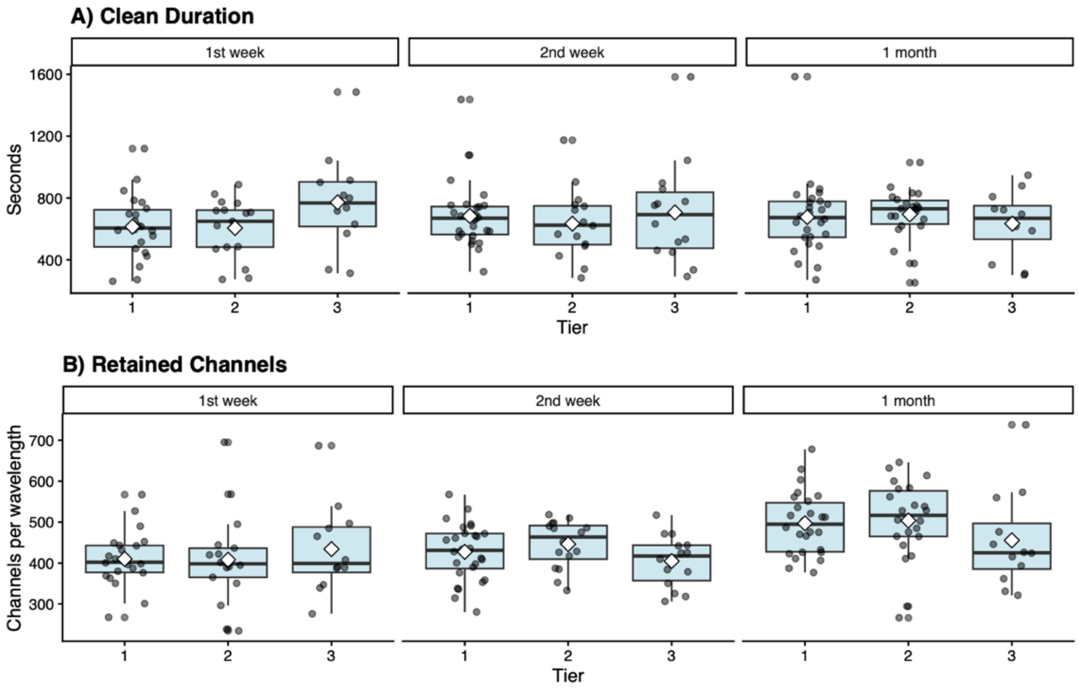


#### e. Seed-based analyses and multiverse thresholding

We select the strongest connections at the individual level, for each of the seeds of interest (Supplementary Figure 11).

*Supplementary Figure 11.* Visualisation of seed location with parcellation drawn on the image. Seed is identified as the red area, while the blue area is that outside our group sensitivity mask

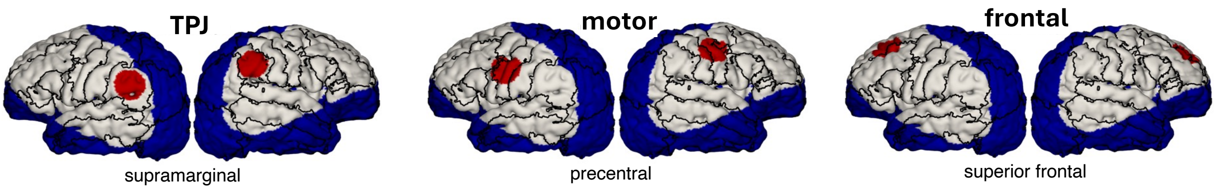


We acknowledge that the choice of a threshold is arbitrary and can influence results. Over-filtering may introduce bias by eliminating relevant data, while under-filtering may allow weak, non-informative connections to remain. Recognize the arbitrary nature of this decision and, inspired by prior research on multiverse analyses [19,20], we ran the analyses under multiple thresholds to assess the robustness of our findings and report on their consistency. More conservative thresholds yield stronger connectivity values (

Supplementary Figure 12), but this might favour short-distance connections (Supplementary Figure 13) that are known to be the strongest at this age [21]. Hence, we compute the value thresholds at different percentiles (bottom 30, 40, 50, 60, 70 in their absolute values) and exclude connectivity values that are weaker than such threshold.

*Supplementary Figure 12.* Distribution of threshold values for different thresholding scenarios, across seeds. Panels on y axis (30 to 70) reflect exclusion of bottom x% connections. Thresholds are computed separately for HbO and HbR, individually for each participant and seed.

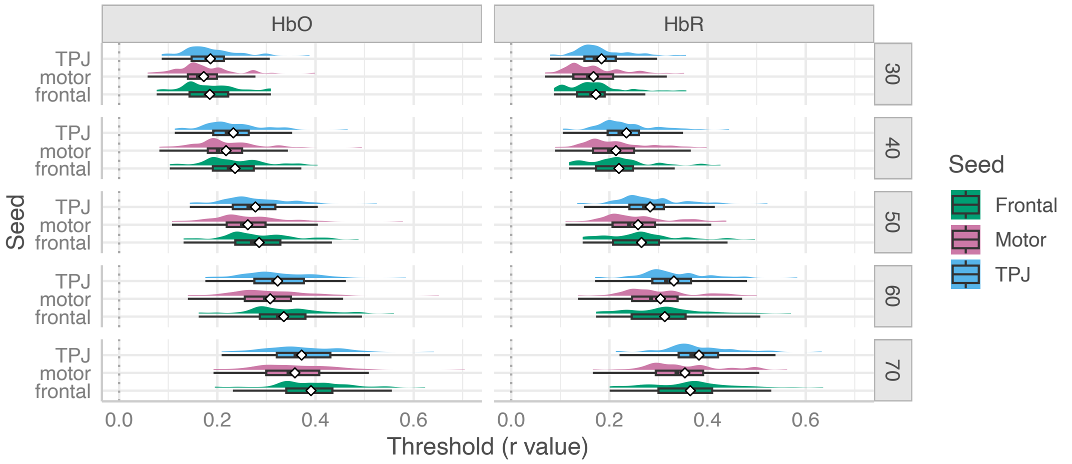


*Supplementary Table 3. Descriptives of threshold (r values) across different thresholding scenarios*

| | | **HbO** | | **HbR** | |
|---|---|---|---|---|---|
| **Exclude bottom %** | **Seed** | **Mean** | **Range** | **Mean** | **Range** |
| **30** | TPJ | 0.186 | 0.087–0.388 | 0.184 | 0.079–0.356 |
| | motor | 0.172 | 0.058–0.399 | 0.168 | 0.069–0.352 |
| | frontal | 0.185 | 0.076–0.309 | 0.172 | 0.087–0.357 |
| **40** | TPJ | 0.233 | 0.113–0.465 | 0.235 | 0.105–0.444 |
| | motor | 0.218 | 0.082–0.494 | 0.213 | 0.089–0.398 |
| | frontal | 0.236 | 0.103–0.404 | 0.219 | 0.117–0.426 |
| **50** | TPJ | 0.278 | 0.144–0.525 | 0.283 | 0.135–0.522 |
| | motor | 0.262 | 0.108–0.578 | 0.258 | 0.111–0.438 |
| | frontal | 0.285 | 0.131–0.488 | 0.265 | 0.145–0.497 |
| **60** | TPJ | 0.323 | 0.176–0.584 | 0.331 | 0.171–0.583 |
| | motor | 0.308 | 0.14–0.651 | 0.304 | 0.136–0.5 |
| | frontal | 0.335 | 0.162–0.56 | 0.313 | 0.173–0.57 |
| **70** | TPJ | 0.372 | 0.209–0.642 | 0.383 | 0.213–0.633 |
| | motor | 0.358 | 0.192–0.704 | 0.354 | 0.166–0.562 |
| | frontal | 0.391 | 0.195–0.624 | 0.365 | 0.2–0.636 |

*Supplementary Figure 13.* Variation in (A) strength and (B) extent by distance category across thresholding levels.

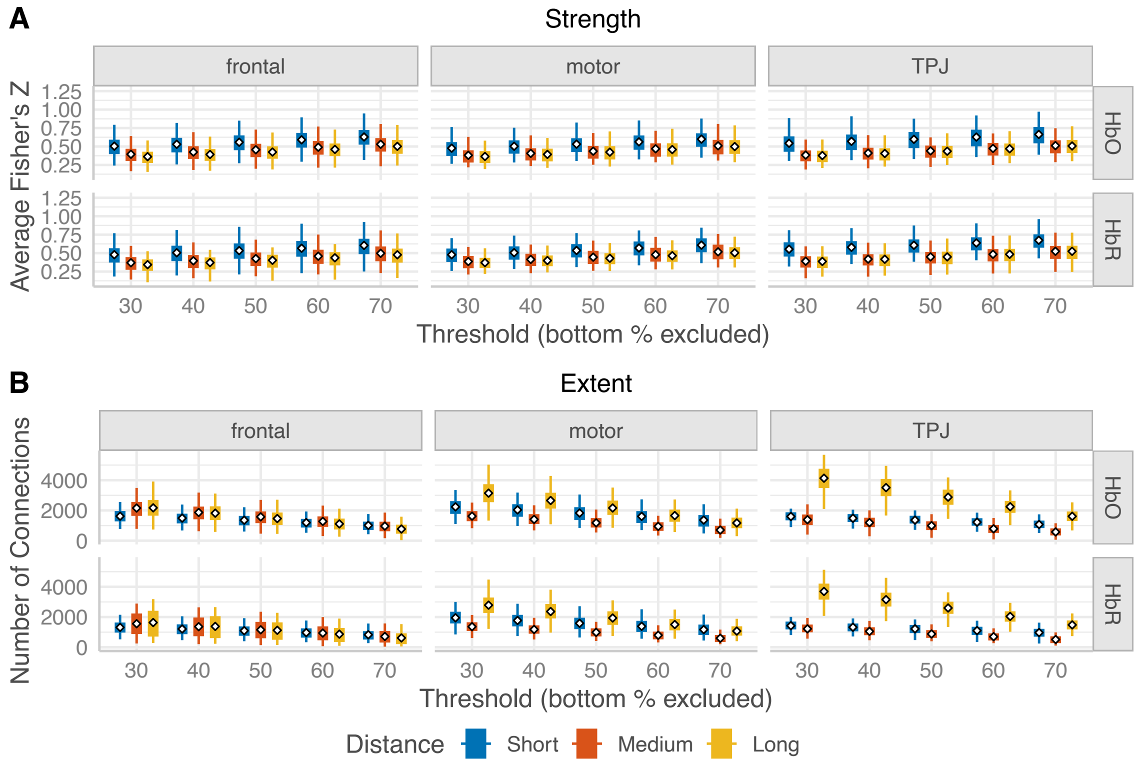

*Supplementary Figure 14.* Group level plots of FC for the different seeds at the 1-month timepoint for both HbO and HbR. Seeds are represented with black dots.

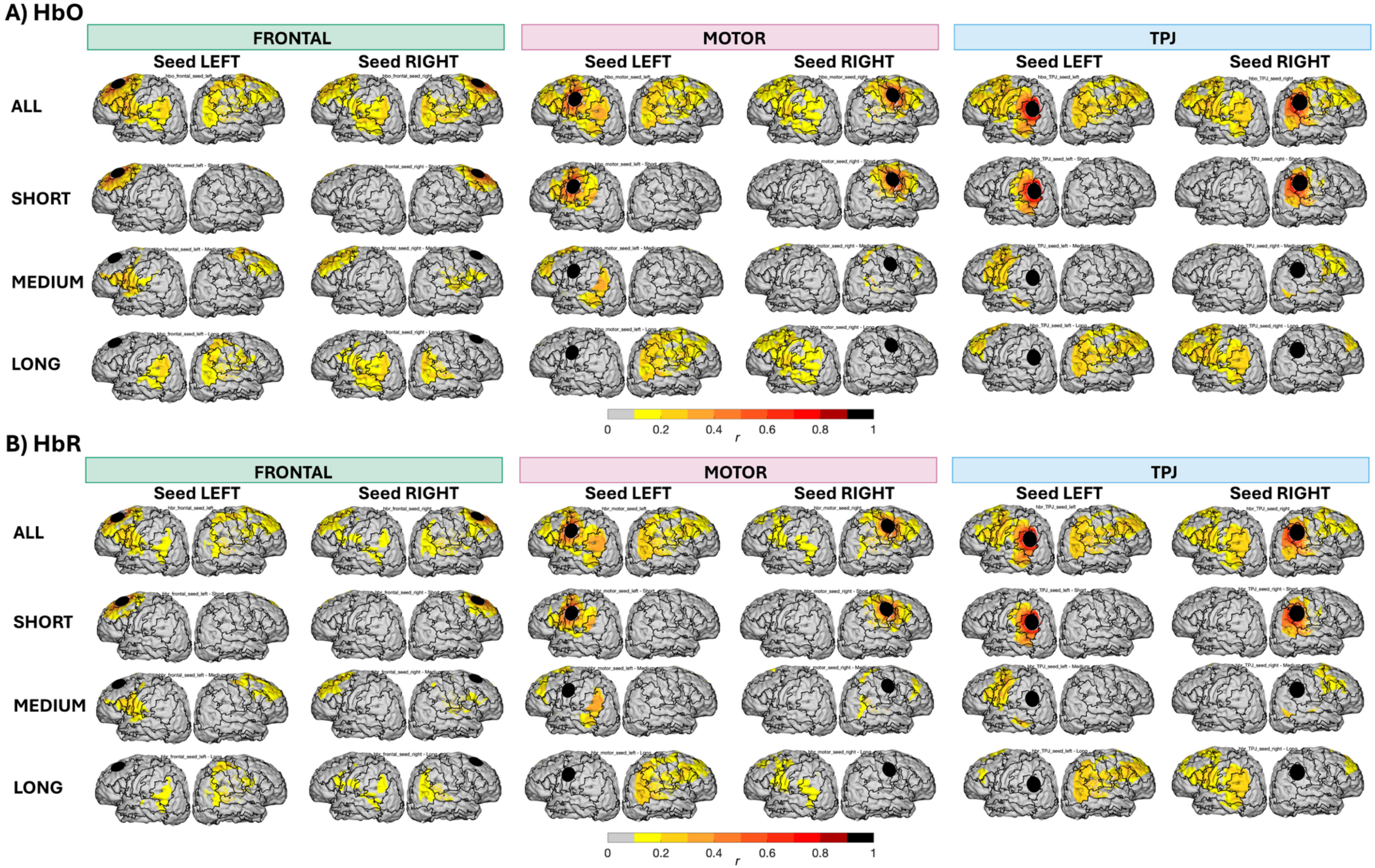

### 4. STATISTICAL ANALYSES

#### a. Model comparisons

*Supplementary Table 4.* Model comparisons for strength and extent, with summary of parameters used for model selections (AIC, AICc and their weights, $R^2$) as well as complete report of statistically significant effects for the selected model (thresholding out bottom 30% - HbO).

| STRENGTH | | | | |
|---|---|---|---|---|
| Model | | AIC (weigths) | AICc (weigths) | R2 (marginal) |
| m0 | (1\|Participant) | -564.8 (<.001) | -564.7 (<.001) | 0.000 |
| m1 | Dyadic Engagement + Affectionate touch + Age + Age at birth + (1\|Participant) | -560.9 (<.001) | -560.6 (<.001) | 0.030 |
| m2 | Dyadic Engagement × Distance + Affectionate touch + Age + Age at birth + (1\|Participant) | -760.4 (<.001) | -759.9 (<.001) | 0.324 |
| m3 | Dyadic Engagement × Distance + Affectionate touch × Distance + Age + Age at birth + (1\|Participant) | -758.2 (<.001) | -757.5 (<.001) | 0.325 |
| m4 | Dyadic Engagement × Distance × Seed + Affectionate touch × Distance + Age + Age at birth + (1\|Participant) | -816.5 (0.994) | -814.8 (0.997) | 0.390 |
| m5 | Dyadic Engagement × Distance × Seed + Affectionate touch × Distance × Seed + Age + Age at birth + (1\|Participant) | -806.3 (0.006) | -803.3 (0.003) | 0.390 |

The selected model's total explanatory power is substantial (conditional R2 = 0.66) and the part related to the fixed effects alone (marginal R2) is of 0.39.
- Dyadic Eng× Distance [Short] × Seed [Frontal] is statistically significant (β = 7.87e-04, 95% CI [2.21e-04, 1.35e-03], t(370) = 2.74, p = 0.007; Std. β = 0.26, 95% CI [-0.06, 0.57])
- Dyadic Eng× Distance [Short] × Seed [TPJ] is statistically significant (β = 1.14e-03, 95% CI [5.75e-04, 1.71e-03], t(370) = 3.97, p < .001; Std. β = 0.07, 95% CI [-0.25, 0.38])

| EXTENT | | | | |
|---|---|---|---|---|
| Model | | AIC (weigths) | AICc (weigths) | R2 (marginal) |
| m0 | (1\|Participant) | 7091.1 (<.001) | 7091.2 (<.001) | 0.000 |
| m1 | Dyadic Engagement + Affectionate touch + Age + Age at birth + (1\|Participant) | 7096.0 (<.001) | 7096.3 (<.001) | 0.008 |
| m2 | Dyadic Engagement × Distance + Affectionate touch + Age + Age at birth+ (1\|Participant) | 6839.5 (<.001) | 6840.0 (<.001) | 0.489 |
| m3 | Dyadic Engagement × Distance + Affectionate touch × Distance + Age + Age at birth + (1\|Participant) | 6841.2 (<.001) | 6841.9 (<.001) | 0.491 |
| m4 | Dyadic Engagement × Distance × Seed + Affectionate touch × Distance + Age + Age at birth + (1\|Participant) | 6551.1 (0.742) | 6552.8 (0.852) | 0.743 |
| m5 | Dyadic Engagement × Distance × Seed + Affectionate touch × Distance × Seed + Age + Age at birth + (1\|Participant) | 6553.2 (0.258) | 6556.3 (0.148) | 0.746 |

The selected model's total explanatory power is substantial (conditional R2 = 0.79) and the part related to the fixed effects alone (marginal R2) is of 0.74.
- Distance [Long] × Affectionate Touch is statistically significant and positive (β = 17.19, 95% CI [5.76, 28.61], t(370) = 2.96, p = 0.003; Std. β = 0.13, 95% CI [-0.06, 0.32])
- Dyadic Eng× Distance [Short] × Seed [Frontal] is statistically significant (β = -6.90, 95% CI [-12.33, -1.47], t(370) = -2.50, p = 0.013; Std. β = 0.11, 95% CI [-0.21, 0.42])
- Dyadic Eng× Distance [Medium] × Seed [Motor] is statistically significant (β = -12.91, 95% CI [-18.34, -7.48], t(370) = -4.67, p < .001; Std. β = -0.05, 95% CI [-0.36, 0.27])
- Dyadic Eng× Distance [Long] × Seed [Motor] is statistically significant (β = 13.00, 95% CI [7.56, 18.43], t(370) = 4.70, p < .001; Std. β = -0.07, 95% CI [-0.38, 0.24])
- Dyadic Eng× Distance [Short] × Seed [TPJ] is statistically significant (β = -11.63, 95% CI [-17.06, -6.19], t(370) = -4.21, p < .001; Std. β = -0.02, 95% CI [-0.33, 0.30])
- Dyadic Eng× Distance [Medium] × Seed [TPJ] is statistically significant(β = -9.65, 95% CI [-15.08, -4.22], t(370) = -3.49, p < .001; Std. β = 0.01, 95% CI [-0.30, 0.33])
- Dyadic Eng× Distance [Long] × Seed [TPJ] is statistically significant (β = 24.88, 95% CI [19.45, 30.31], t(370) = 9.01, p < .001; Std. β = -0.07, 95% CI [-0.38, 0.25])

*Supplementary Figure 15.* Changes in Strength and Extent of connectivity across timepoints, quantified via individual-level slopes (for strength: AvgZ ~ GW; for extent: NumConn ~ GW) extracted for each combination of seed location and distance.

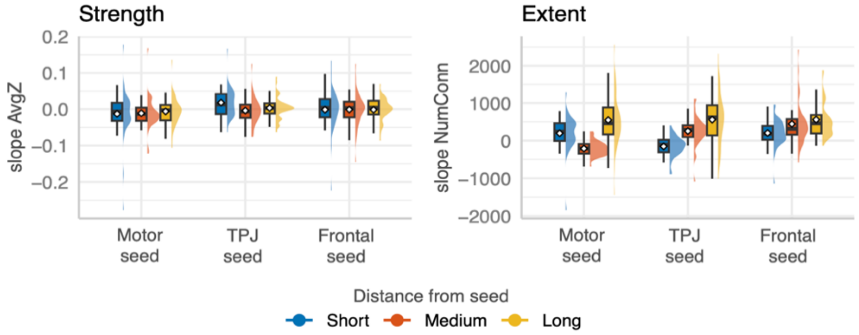


We examined longitudinal changes in the extent of FC across the first month of life. We focused on the extent metric as preliminary analyses indicated that longitudinal effects were more pronounced for connection extent than for strength (Supplementary Figure 15); therefore, extent was selected as the primary outcome for longitudinal modelling.

*Supplementary Table 5.* Model comparisons for longitudinal change in FC extent, with summary of parameters used for model selections (AIC, AICc and their weights, R2) as well as report of statistically significant effects for the selected model (thresholding out bottom 30% - HbO).

| LONGITUDINAL CHANGE IN FUNCTIONAL CONNECTIVITY EXTENT | | | |
|---|---|---|---|
| Model | AIC (weigths) | AICc (weigths) | R2 (marginal) |
| m0 (1\|Participant) | 5127.1 (<.001) | 5127.2 (<.001) | 0.000 |
| m1 Distance + Seed + Sex + Age at birth + (1\|Participant) | 5053.3 (<.001) | 5053.9 (<.001) | 0.182 |
| m2 Distance × Seed + Sex + Age at birth + (1\|Participant) | 5011.3 (>.999) | 5012.4 (>.999) | 0.256 |

The selected model's total explanatory power is substantial (conditional R2 = 0.53) and the part related to the fixed effects alone (marginal R2) is of 0.26. Post-hoc analyses with FDR correction for multiple comparisons revealed the following pattern of estimated marginal means:

| | | |
|---|---|---|
| Motor seed | Short | est = 183, SE = 82.8, t(142) = 2.21, 95% CI [-50.01,416.17], p = 0.0322 |
| | Medium | est = -224, SE = 82.8, t(142) = -2.7, 95% CI [-456.94,9.23], p = 0.0115 |
| | Long | est = 532, SE = 82.8, t(142) = 6.43, 95% CI [298.76,764.93], p <.0001 |
| TPJ seed | Short | est = -167, SE = 82.8, t(142) = -2.02, 95% CI [-400.22,65.95], p = 0.0453 |
| | Medium | est = 240, SE = 82.8, t(142) = 2.9, 95% CI [7.01,473.18], p = 0.0078 |
| | Long | est = 542, SE = 82.8, t(142) = 6.55, 95% CI [309.22,775.39], p <.0001 |
| Frontal seed | Short | est = 189, SE = 82.8, t(142) = 2.28, 95% CI [-44.13,422.04], p = 0.0308 |
| | Medium | est = 430, SE = 82.8, t(142) = 5.19, 95% CI [196.42,662.59], p <.0001 |
| | Long | est = 543, SE = 82.8, t(142) = 6.56, 95% CI [310.24,776.41], p <.0001 |

| BEHAVIOURS AND LONGITUDINAL CHANGE IN FUNCTIONAL CONNECTIVITY EXTENT | | | |
|---|---|---|---|
| Model | AIC (weigths) | AICc (weigths) | R2 (marginal) |
| m0 (1\|Participant) | 5127.1 (<.001) | 5127.2 (<.001) | 0.000 |
| m1 Dyadic Eng + Age at birth + (1\|Participant) | 5129.9 (<.001) | 5130.1 (<.001) | 0.011 |
| m2 Dyadic Eng × Distance + Age at birth + (1\|Participant) | 5070.4 (<.001) | 5071.0 (<.001) | 0.141 |
| m3 Dyadic Eng × Distance + Dyadic Eng × Seed + Age at birth + (1\|Participant) | 5067.0 (<.001) | 5067.5 (<.001) | 0.147 |
| m4 Dyadic Eng × Distance × Seed + Age at birth + (1\|Participant) | 5026.2 (>.999) | 5029.2 (>.999) | 0.241 |

The model's total explanatory power is substantial (conditional R2 = 0.53) and the part related to the fixed effects alone (marginal R2) is of 0.24. Within this model, the interaction of Distance × Seed was significant (same pattern as in model set above).

| | Model | AIC (weigths) | AICc (weigths) | R2 (marginal) |
|---|---|---|---|---|
| m0 | (1\|Participant) | 5127.1 (<.001) | 5127.2 (<.001) | 0.000 |
| m1 | Affectionate Touch + Age at birth + (1\|Participant) | 5128.2 (<.001) | 5128.4 (<.001) | 0.026 |
| m2 | Affectionate Touch × Distance + Age at birth + (1\|Participant) | 5090.6 (<.001) | 5090.9 (<.001) | 0.109 |
| m3 | Affectionate Touch × Distance + Affectionate Touch × Seed + Age at birth + (1\|Participant) | 5081.1 (<.001) | 5081.7 (<.001) | 0.134 |
| m4 | Affectionate Touch × Distance × Seed + Age at birth + (1\|Participant) | 5015.4 (>.999) | 5018.4 (>.999) | 0.268 |

The model's total explanatory power is substantial (conditional R2 = 0.54) and the part related to the fixed effects alone (marginal R2) is of 0.27.

- The effect of Distance [Short] × Seed [Motor] is statistically significant (β = -498.43, 95% CI [-836.92, -159.94], t(312) = -2.90, p = 0.004; Std. β = -0.63, 95% CI [-0.95, -0.32])
- The effect of Distance [Medium] × Seed [Motor] seed is statistically significant (β = -856.48, 95% CI [-1194.97, -517.99], t(312) = -4.98, p < .001; Std. β = -1.35, 95% CI [-1.67, -1.03])
- The effect of Distance [Short] × Seed [TPJ] is statistically significant (β = -793.14, 95% CI [-1131.63, -454.65], t(312) = -4.61, p < .001; Std. β = -1.25, 95% CI [-1.57, -0.93])
- The effect of Distance [Medium] × Seed [TPJ] is statistically significant (β = -571.38, 95% CI [-909.86, -232.89], t(312) = -3.32, p = 0.001; Std. β = -0.53, 95% CI [-0.85, -0.22])
- The effect of Distance [Short] × Seed [Frontal] is statistically significant (β = -567.67, 95% CI [-906.16, -229.18], t(312) = -3.30, p = 0.001; Std. β = -0.62, 95% CI [-0.94, -0.31])
- The effect of Distance [Medium] × Seed [Frontal] is statistically significant (β = -433.15, 95% CI [-771.64, -94.66], t(312) = -2.52, p = 0.012; Std. β = -0.20, 95% CI [-0.52, 0.12])
- The effect of Affectionate Touch × Distance [Long] × Seed [TPJ] is statistically significant (β = 7.27, 95% CI [0.62, 13.92], t(312) = 2.15, p = 0.032; Std. β = 0.31, 95% CI [0.03,0.59])
- The effect of Affectionate Touch × Distance [Medium] × Seed [Frontal] is statistically significant (β = 6.90, 95% CI [0.26, 13.55], t(312) = 2.04, p = 0.042; Std. β = 0.29, 95% CI [0.01, 0.58])

#### b. Consistency of results across thresholding scenarios and chromophores

Supplementary Figure 16. Consistency of results (1 month) across chromophores and thresholds

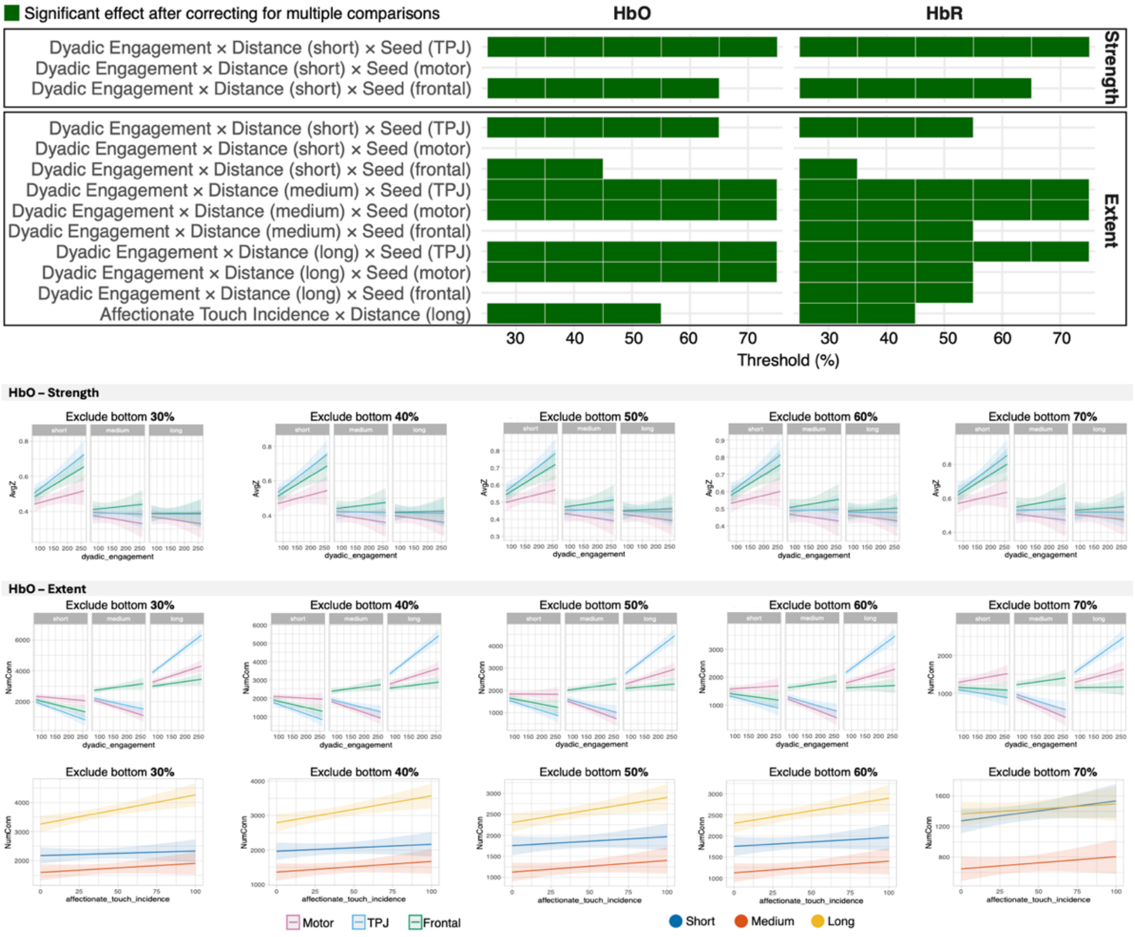

#### c. Variance partitioning analyses to check influences on dyadic engagement composite

To assess whether variability in the dyadic engagement composite was primarily driven by infant behaviour, parent behaviour, or changes in FC extent, we conducted a variance decomposition of the composite score. Since dyadic engagement was computed as the sum of its behavioural components (infant-to-parent gaze, parent-to-infant gaze, and affectionate touch occurring during infant attention), we decomposed the total variance of the composite using the algebraic identity $Var(X+Y+Z) = Var(X) + Var(Y) + Var(Z) + 2Cov(X,Y) + 2Cov(X,Z) + 2Cov(Y,Z)$, which allowed us to quantify the unique variance contribution of each component as well as their shared covariance terms. To assess the independent relationship between change in functional connectivity extent and the composite score, we additionally fitted a simple linear regression of dyadic engagement on connectivity slopes separately for each seed × distance combination (9 models total), controlling for no other variables given the orthogonality of connectivity to the behavioural components.

Variance decomposition of the dyadic engagement composite revealed infant-to-parent gaze accounted for the largest unique proportion of variance (30.76%). However, additional variance was explained by affectionate touch occurring during infant attention (20.96%) and parent-to-infant gaze (3.78%). The remaining variance (~44.5%) was attributable to covariance between components, most notably between infant-to-parent gaze and affectionate touch (32.61%), reflecting the tendency for these behaviours to co-occur. In contrast, change in functional connectivity extent showed no meaningful relationship with the dyadic engagement composite across any seed region or distance level ($R^2 < 2\%$ in all cases). These findings indicate that, while infant attention has notable weight in our dyadic engagement composite, a substantial proportion of variance also reflects parental behaviour, supporting the interpretation of the measure as capturing dyadic rather than purely infant-driven engagement. Detailed percentages of variance explained by each component are reported in (Supplementary Table 6 and

Supplementary Table 7).

*Supplementary Table 6.* Proportions of variance in dyadic engagement composite explained by each of infant to parent gaze, parent to infant gaze and affectionate touch to boost.

| Component | Pct of Total Variance (%) |
|---|---|
| Infant-to-parent gaze | 30.76 |
| Parent-to-infant gaze | 3.78 |
| Affectionate touch (during infant attention) | 20.96 |
| Cov(infant-to-parent; parent-to-infant) | 8.17 |
| Cov(infant-to-parent; touch) | 32.61 |
| Cov(parent-to-infant; touch) | 3.72 |

*Supplementary Table 7.* Variance in dyadic engagement composite explained by longitudinal change in functional connectivity extent

| Seed | Distance | $R^2$ |
|---|---|---|
| Motor | Short | 0.00 |
| | Medium | 1.12 |
| | Long | 0.36 |
| TPJ | Short | 0.01 |
| | Medium | 1.89 |
| | Long | 0.00 |
| Frontal | Short | 1.50 |
| | Medium | 0.66 |
| | Long | 1.20 |

#### d. Ruling out socioeconomic status as a confound

To examine whether the observed associations between functional connectivity and parent-infant interaction can be explained by underlying socioeconomic factors, we performed two different analyses controlling for subjective social status and for household income.

Subjective social status was assessed using the MacArthur Social Ladder scale [22], which captures individuals' perceived position within the social hierarchy. This measure has been shown to relate not only to conventional SES indicators (i.e. income) but also psychosocial well-being in general [23] and may therefore be relevant for explaining variation in caregiving behaviours. While it might be related to income, it is not solely impacted by that, as depicted in Supplementary Figure 17.

Household income was included as an additional SES proxy. Income was measured as a categorical variable with the following categories: "< £1,400", "£1,400-£2,000", "£2,000-£2,600", "£2,600-£3,600", "£3,600-£4,600", "£4,600-£5,600", "£5,600-£6,600", "£6,600-£8,000", "£8,000-£10,000", "> £10,000", and "Do not wish to answer". Due to the ordered nature of the income scale and the need to use income as part of the correlational analysis, the response was coded onto an ordinal numeric scale from 1 to 10, with those answering "Do not wish to answer" excluded. While this imposes a simplified linear structure on categorical data, it provides a pragmatic approximation of income variation.

SES data were not available for all participants, as some individuals opted not to provide responses for income, subjective social status, or both. Consequently, analyses including these variables were conducted on a reduced subsample.

As shown in Supplementary Figure 17, the range of income levels and subjective social ladder scores, indicates some diversity in socioeconomic and psychosocial context. However, this diversity did not translate to diversity in education levels, as most parents had an undergraduate degree as their highest qualification, and a significant proportion holds postgraduate degrees (Supplementary Figure 17C). Therefore, education was not used in the analyses because of the low diversity and unbalanced distribution in categories, although it is presented descriptively. For providing additional context on our sample composition, ethnicity is also described (Supplementary Figure 17D).

*Supplementary Figure 17.* Visualisation of SES and demographic information in our sample. **A)** Income category counts; **B)** social ladders scores across income categories; **C)** Education of mothers and partners; **D)** Ethnicity of mothers and babies.

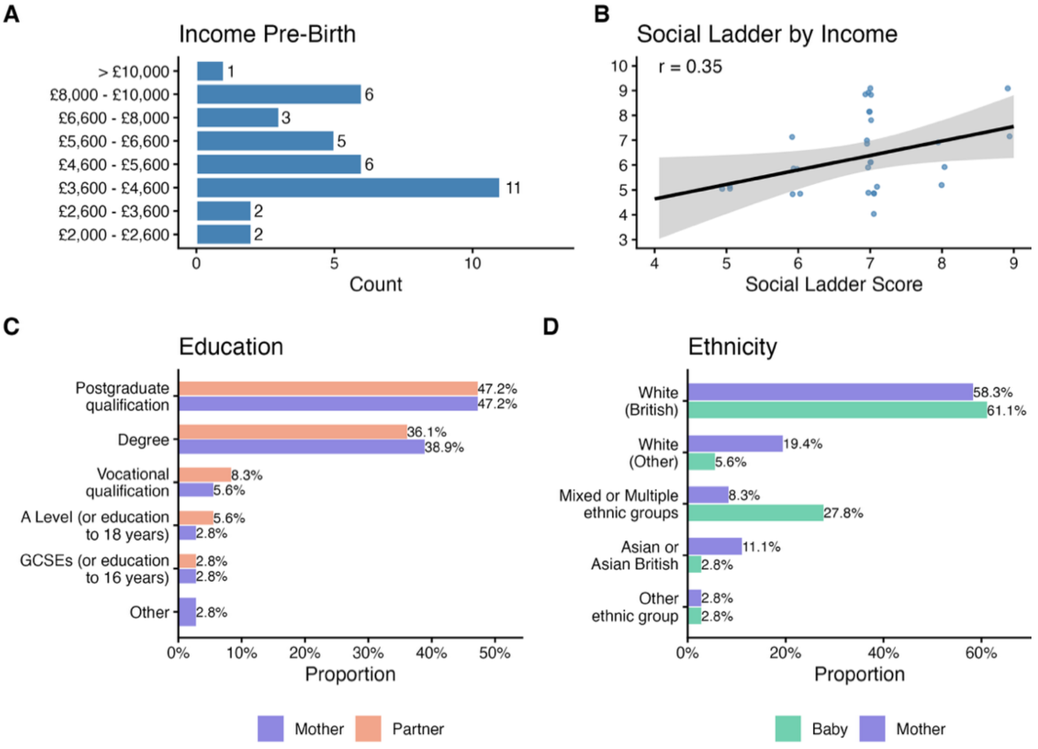


Overall, SES in the present study should be interpreted as a broad proxy for psychosocial context rather than a precise index of material resources alone. In particular, subjective social status may capture dimensions such as perceived stability, stress, and social support, which are plausibly more proximal to caregiving behaviours, including touch.

**Cross-sectional analyses (1 month)**

For the cross-sectional models, we compared our selected model against two SES-adjusted variants, which included social ladder scores as a main effect or both interactions between social ladder and parenting behaviours. The same approach was taken to assess the influence of Income, in a separate set of models.

For **FC strength (AvgZ)**, model fit indices strongly supported the original model over both SES-adjusted variants, and the addition of both Social Ladder and Household Income had negligible effects on marginal $R^2$, suggesting that SES (in terms of both subjective social status and income) explains little variance in FC strength (Supplementary Table 8 and Supplementary Table 9).

*Supplementary Table 8* Model performance of selected model with and without Social Ladder for Strength

| PREDICTORS | AIC | $R^2$ (marg) |
|---|---|---|
| Dyadic Engagement × Distance × Seed + Affectionate touch × Distance + Age + Age at birth + (1\|Participant) | -816.45 | 0.39 |
| Dyadic Engagement × Distance × Seed + Affectionate touch × Distance + Age + Age at birth + Social Ladder + (1\|Participant) | -749.84 | 0.41 |
| Dyadic Engagement × Distance × Seed + Affectionate touch × Distance + Age + Age at birth + Dyadic Engagement × Social Ladder + Affectionate Touch × Social Ladder + (1\|Participant) | -750.52 | 0.42 |

*Supplementary Table 9* Model performance of selected model with and without Household Income for Strength

| PREDICTORS | AIC | $R^2$ (marg) |
|---|---|---|

| | | |
|---|---|---|
| Dyadic Engagement × Distance × Seed + Affectionate touch × Distance + Age + Age at birth + (1\|Participant) | -810.5 | 0.39 |
| Dyadic Engagement × Distance × Seed + Affectionate touch × Distance + Age + Age at birth + Household Income + (1\|Participant) | -677.3 | 0.41 |
| Dyadic Engagement × Distance × Seed + Affectionate touch × Distance + Age + Age at birth + Dyadic Engagement × Household Income + Affectionate Touch × Household Income + (1\|Participant) | -677.6 | 0.42 |

For **FC extent (NumConn)**, the SES-adjusted models showed somewhat better fit (Supplementary Table 10, Supplementary Table 11), suggesting that SES (in terms of both subjective social status and income) does explain some variance in the number of functional connections. Nevertheless, the associations found in the initial model were replicated, with coefficients changing only minimally. Importantly, social ladder did not emerge as a significant predictor (β = 11.61, SE = 104.21, t(33) = 0.11, p = 0.911) and neither did Household income (β = -44.56, SE = 56.87, t(30) = 0.78, p = 0.439) suggesting that while they help explaining more variance neither constitute a discernible effect. The pattern of significant effects is generally consistent across models with and without SES predictors (Supplementary Table 12).

*Supplementary Table 10.* Model performance of selected model with and without Social Ladder for Extent

| PREDICTORS | AIC | $R^2$ (marg) |
|---|---|---|
| Dyadic Engagement × Distance × Seed + Affectionate touch × Distance + Age + Age at birth + (1\|Participant) | 6551.11 | 0.743 |
| Dyadic Engagement × Distance × Seed + Affectionate touch × Distance + Age + Age at birth + Social Ladder + (1\|Participant) | 5951.23 | 0.738 |
| Dyadic Engagement × Distance × Seed + Affectionate touch × Distance + Age + Age at birth + Dyadic Engagement × Social Ladder + Affectionate Touch × Social Ladder + (1\|Participant) | 5951.54 | 0.740 |

*Supplementary Table 11.* Model performance of selected model with and without Household Income for Extent

| PREDICTORS | AIC | $R^2$ (marg) |
|---|---|---|
| Dyadic Engagement × Distance × Seed + Affectionate touch × Distance + Age + Age at birth + (1\|Participant) | 6582.7 | 0.726 |
| Dyadic Engagement × Distance × Seed + Affectionate touch × Distance + Age + Age at birth + Household Income + (1\|Participant) | 5521.3 | 0.733 |
| Dyadic Engagement × Distance × Seed + Affectionate touch × Distance + Age + Age at birth + Dyadic Engagement × Household Income + Affectionate Touch × Household Income + (1\|Participant) | 5522.0 | 0.733 |

*Supplementary Table 12. Summary of significant effects across models with and without social ladder*

| | | |
|---|---|---|
| **Affectionate touch × Distance** | | |
| Long | Original | β = 1.72 × 10¹, SE = 5.81, t(95.32) = 2.96, p = .004 |
| | *with Social Ladder* | β = 1.66 × 10¹, SE = 6.49, t(73.14) = 2.57, p = .012 |
| | *with Household Income* | β = 2.29 × 10¹, SE = 5.9, t(81.57) = 3.88, p < .001 |
| **Dyadic Engagement × Distance × Seed** | | |
| Motor (Medium) | Original | β = -1.29 × 10¹, SE = 2.76, t(65.2) = -4.67, p < .001 |
| | *with Social Ladder* | β = -1.18 × 10¹, SE = 3.1, t(53.44) = -3.82, p < .001 |
| | *with Household Income* | β = -8.78, SE = 1.96, t(70.96) = -4.47, p < .001 |
| Motor (Long) | Original | β = 1.3 × 10¹, SE = 2.76, t(65.2) = 4.7, p < .001 |
| | *with Social Ladder* | β = 1.33 × 10¹, SE = 3.1, t(53.44) = 4.28, p < .001 |
| | *with Household Income* | β = 9.62, SE = 1.96, t(70.96) = 4.9, p < .001 |
| | Original | β = -1.16 × 10¹, SE = 2.76, t(65.2) = -4.21, p < .001 |

| | | |
|---|---|---|
| TPJ (Short) | *with Social Ladder* | $\beta = -1.08 \times 10^1$, SE = 3.1, t(53.44) = -3.47, p = .001 |
| | *with Household Income* | β = -7.79, SE = 1.96, t(70.96) = -3.97, p < .001 |
| TPJ (Medium) | Original | β = -9.65, SE = 2.76, t(65.2) = -3.49, p < .001 |
| | *with Social Ladder* | β = -8.58, SE = 3.1, t(53.44) = -2.76, p = .008 |
| | *with Household Income* | β = -5.95, SE = 1.96, t(70.96) = -3.03, p = 0.003 |
| TPJ (Long) | Original | $\beta = 2.49 \times 10^1$, SE = 2.76, t(65.2) = 9.01, p < .001 |
| | *with Social Ladder* | $\beta = 2.51 \times 10^1$, SE = 3.1, t(53.44) = 8.09, p < .001 |
| | *with Household Income* | $\beta = 1.81 \times 10^1$, SE = 1.96, t(70.96) = 9.25, p < .001 |
| Frontal (Short) | Original | β = -6.9, SE = 2.76, t(65.2) = -2.5, p = .015 |
| | *with Social Ladder* | β = -4.2, SE = 2.06, t(64.72) = -2.04, p = 0.045 |
| | *with Household Income* | β = -4.05, SE = 1.96, t(70.96) = -2.06, p = 0.043 |

Overall, smaller standard error and larger t-value were found in the original models (without social ladder or household income), which is expected given the increased degrees of freedom reflecting the larger sample size when SES variables, with missing data in some cases, are not included in the analysis. Importantly, the associations of interest remained significant after adjustment for both subjective social status and household income, suggesting that while they help explaining a slightly more variance overall (around +1%) these SES measures do not account for the observed relationships between parenting behaviour and infant functional connectivity.

**Longitudinal analyses**

For the longitudinal models we took the same approach detailed above. Specifically, we compared our selected model against two SES-adjusted variants, which included social ladder scores and income (in separate model sets) as a main effect or both interactions between social ladder and parenting behaviours.

The inclusion of SES as a covariate improved overall model fit as indicated by lower AIC (For Dyadic Engagement see Supplementary Table 13 and Supplementary Table 14, For Affectionate touch see Supplementary Table 15 and Supplementary Table 16), suggesting that SES explains additional variance in longitudinal change in connectivity extent. However, neither social ladder nor household income showed significant main effects, and there was no evidence that SES moderated associations between parent behaviour and longitudinal change in FC.

Consistent with the main analyses, affectionate touch continued to show seed- and distance-specific associations with longitudinal change in connectivity (e.g., TPJ long-range and frontal medium-range effects, see Supplementary Table 17), whereas dyadic engagement remained unrelated to connectivity change across all models. While this pattern of significant effects is generally consistent across models with and without SES predictors (Supplementary Table 12), some attenuation of the effect was observed in frontal connectivity, which became only marginally significant. A caveat to this though is that sample size reduces when implementing this SES adjustment due to incomplete SES data (from n = 37 participants included for longitudinal analyses, sample size becomes n = 33 with Social Ladder and n = 34 with Household Income), which might have decreased power in this case.

*Supplementary Table 13* Model performance of selected model for Dyadic Engagement with and without **Social Ladder** for Longitudinal Change in Number of Connections (slopes of extent)

| PREDICTORS | AIC (weight) | $R^2$ (marg) |
|---|---|---|
| Dyadic Engagement × Distance × Seed + Age at birth + (1\|Participant) | 5026.2 (<.001) | 0.241 |
| Dyadic Engagement × Distance × Seed + Age at birth + Social Ladder + (1\|Participant) | 4437.0 (0.512) | 0.269 |
| Dyadic Engagement × Distance × Seed + Age at birth + Dyadic Engagement × Social Ladder + (1\|Participant) | 4437.1 (0.488) | 0.268 |

*Supplementary Table 14* Model performance of selected model for Dyadic Engagement with and without **Household Income** for Longitudinal Change in Number of Connections (slopes of extent)

| PREDICTORS | AIC (weight) | $R^2$ (marg) |
|---|---|---|
| Dyadic Engagement × Distance × Seed + Age at birth + (1\|Participant) | 5026.2 (<.001) | 0.241 |
| Dyadic Engagement × Distance × Seed + Age at birth + Household Income + (1\|Participant) | 4560.7 (0.519) | 0.284 |
| Dyadic Engagement × Distance × Seed + Age at birth + Dyadic Engagement × Household Income + (1\|Participant) | 4560.8 (0.481) | 0.283 |

*Supplementary Table 15* Model performance of selected model for Affectionate Touch with and without **Social Ladder** for Longitudinal Change in Number of Connections (slopes of extent)

| PREDICTORS | AIC (weight) | $R^2$ (marg) |
|---|---|---|
| Affectionate Touch × Distance × Seed + Age at birth + (1\|Participant) | 5015.4 (<.001) | 0.268 |
| Affectionate Touch × Distance × Seed + Age at birth + Social Ladder + (1\|Participant) | 4428.7 (0.552) | 0.287 |
| Affectionate Touch × Distance × Seed + Age at birth + Affectionate Touch × Social Ladder + (1\|Participant) | 4429.1 (0.448) | 0.283 |

*Supplementary Table 16* Model performance of selected model for Affectionate Touch with and without **Household Income** for Longitudinal Change in Number of Connections (slopes of extent)

| PREDICTORS | AIC (weight) | $R^2$ (marg) |
|---|---|---|
| Affectionate Touch × Distance × Seed + Age at birth + (1\|Participant) | 5015.4 (<.001) | 0.268 |
| Affectionate Touch × Distance × Seed + Age at birth + Household Income + (1\|Participant) | 4551.9 (0.494) | 0.302 |
| Affectionate Touch × Distance × Seed + Age at birth + Affectionate Touch × Household Income + (1\|Participant) | 4551.8 (0.506) | 0.302 |

*Supplementary Table 17. Summary of significant effects across models with and without social ladder for Affectionate Touch predicting Longitudinal Change in FC. It is relevant to note that sample size varies across models when implementing this SES adjustment due to incomplete SES data (from n = 37 participants included for longitudinal analyses, sample size becomes n = 33 with Social Ladder and n = 34 with Household Income).*

| **Affectionate Touch × Distance × Seed** | | |
|---|---|---|
| TPJ (Long) | Original | β = 7.27, SE = 3.38, t(143.85) = 2.15, p = 0.033 |
| | *with Social Ladder* | β = 6.87, SE = 3.43, t(102.87) = 2, p = 0.048 |
| | *with Household Income* | β = 6.8, SE = 3.19, t(124.77) = 2.13, p = 0.035 |
| Frontal (Medium) | Original | β = 6.9, SE = 3.38, t(143.85) = 2.04, p = 0.043 |
| | *with Social Ladder* | β = 6.11, SE = 3.43, t(102.87) = 1.78, p = 0.077 |
| | *with Household Income* | β = 6.01, SE = 3.19, t(124.77) = 1.88, p = 0.0620 |

### 5. REFERENCES

1. Stack, D. M. & Jean, A. D. L. Communicating through touch: Touching during parent-infant interactions. in *The handbook of touch: Neuroscience, behavioral, and health perspectives* 273–298 (Springer Publishing Company, New York, NY, US, 2011).
2. Crucianelli, L. *et al.* The mindedness of maternal touch: An investigation of maternal mind-mindedness and mother-infant touch interactions. *Developmental Cognitive Neuroscience* **35**, 47–56 (2019).
3. Murray, L., Fiori-Cowley, A., Hooper, R. & Cooper, P. The impact of postnatal depression and associated adversity on early mother-infant interactions and later infant outcome. *Child Dev* **67**, 2512–2526 (1996).
4. Ilyka, D., Johnson, M. H. & Lloyd-Fox, S. Infant social interactions and brain development: A systematic review. *Neuroscience and Biobehavioral Reviews* **130**, 448–469 (2021).
5. Gamer, M., Lemon, J., Fellows, I. & Singh, P. irr: Various coefficients of interrater reliability and agreement (Version 0.84. 1). *R Foundation for Statistical Computing* (2019).

6. Hallgren, K. A. Computing Inter-Rater Reliability for Observational Data: An Overview and Tutorial. *Tutor Quant Methods Psychol* **8**, 23–34 (2012).
7. Huppert, T. J., Diamond, S. G., Franceschini, M. A. & Boas, D. A. HomER: a review of time-series analysis methods for near-infrared spectroscopy of the brain. *Appl. Opt., AO* **48**, D280–D298 (2009).
8. Schweiger, M. & Arridge, S. R. The Toast++ software suite for forward and inverse modeling in optical tomography. *JBO* **19**, 040801 (2014).
9. Uchitel, J. *et al.* Cot-side imaging of functional connectivity in the developing brain during sleep using wearable high-density diffuse optical tomography. *Neuroimage* **265**, 119784 (2023).
10. Vidal-Rosas, E. E. *et al.* Evaluating a new generation of wearable high-density diffuse optical tomography technology via retinotopic mapping of the adult visual cortex. *Neurophotonics* **8**, 025002 (2021).
11. Caballero-Gaudes, C. & Reynolds, R. C. Methods for cleaning the BOLD fMRI signal. *NeuroImage* **154**, 128–149 (2017).
12. Blanco, B. *et al.* Group-level cortical functional connectivity patterns using fNIRS: assessing the effect of bilingualism in young infants. *Neurophotonics* **8**, 025011 (2021).
13. Collins-Jones, L. H. *et al.* Longitudinal infant fNIRS channel-space analyses are robust to variability parameters at the group-level: An image reconstruction investigation. *NeuroImage* **237**, (2021).
14. Alexander, B. *et al.* Desikan-Killiany-Tourville Atlas Compatible Version of M-CRIB Neonatal Parcellated Whole Brain Atlas: The M-CRIB 2.0. *Front Neurosci* **13**, 34 (2019).
15. Makropoulos, A. *et al.* The developing human connectome project: A minimal processing pipeline for neonatal cortical surface reconstruction. *NeuroImage* **173**, 88–112 (2018).
16. Avants, B. B., Epstein, C. L., Grossman, M. & Gee, J. C. Symmetric diffeomorphic image registration with cross-correlation: Evaluating automated labeling of elderly and neurodegenerative brain. *Medical Image Analysis* **12**, 26–41 (2008).
17. Collins-Jones, L. H. *et al.* Whole-head high-density diffuse optical tomography to map infant audio-visual responses to social and non-social stimuli. *Imaging Neuroscience* **2**, imag–2–00244 (2024).

18. Speh, E. *et al.* NeuroDOT: a Matlab and Python toolbox for diffuse optical tomographic brain mapping. in *Optical Tomography and Spectroscopy of Tissue XVI* vol. 13314 163–167 (SPIE, 2025).
19. Calignano, G., Girardi, P. & Altoè, G. First steps into the pupillometry multiverse of developmental science. *Behav Res* **56**, 3346–3365 (2024).
20. Montuori, C., Gambarota, F., Altoé, G. & Arfé, B. The cognitive effects of computational thinking: A systematic review and meta-analytic study. *Computers & Education* **210**, 104961 (2024).
21. Sylvester, C. M. *et al.* Network-specific selectivity of functional connections in the neonatal brain. *Cereb Cortex* **33**, 2200–2214 (2022).
22. Adler, N. E., Epel, E. S., Castellazzo, G. & Ickovics, J. R. Relationship of subjective and objective social status with psychological and physiological functioning: Preliminary data in healthy, White women. *Health Psychology* **19**, 586–592 (2000).
23. Zell, E., Strickhouser, J. E. & Krizan, Z. Subjective social status and health: A meta-analysis of community and society ladders. *Health Psychology* **37**, 979–987 (2018).
